# A Latent Activated Olfactory Stem Cell State Revealed by Single-Cell Transcriptomic and Epigenomic Profiling

**DOI:** 10.1101/2023.10.26.564041

**Authors:** Koen Van den Berge, Dana Bakalar, Hsin-Jung Chou, Divya Kunda, Davide Risso, Kelly Street, Elizabeth Purdom, Sandrine Dudoit, John Ngai, Whitney Heavner

## Abstract

The olfactory epithelium is one of the few regions of the nervous system that sustains neurogenesis throughout life. Its experimental accessibility makes it especially tractable for studying molecular mechanisms that drive neural regeneration in response to injury. In this study, we used single-cell sequencing to identify the transcriptional cascades and epigenetic processes involved in determining olfactory epithelial stem cell fate during injury-induced regeneration. By combining gene expression and accessible chromatin profiles of individual lineage-traced olfactory stem cells, we identified transcriptional heterogeneity among activated stem cells at a stage when cell fates are being specified. We further identified a subset of resting cells that appears poised for activation, characterized by accessible chromatin around wound response and lineage-specific genes prior to their later expression in response to injury. Together these results provide evidence for a latent activated stem cell state, in which a subset of quiescent olfactory epithelial stem cells are epigenetically primed to support injury-induced regeneration.

## Introduction

The generation of cellular diversity in the nervous system requires specification of discrete cell lineages from multipotent neural progenitor cells. Neural tissues that are uniquely capable of regenerating multiple lineages throughout adulthood can undergo frequent cellular turnover under both homeostatic conditions and in response to injury. Regenerative capacity in the nervous system therefore requires neural progenitor cell maintenance, the ability to specify and generate multiple cell lineages, and the ability to respond to acute injury.

The olfactory epithelium (OE) is one of the few sites in the nervous system that supports active neurogenesis throughout life (reviewed in (Denans, Baek, and Piotrowski 2019)). The OE divides these capacities between two different progenitor cell populations. Under normal homeostatic conditions, the differentiation of globose basal cells (GBCs), the actively proliferating neural progenitor cells in the OE stem cell niche, sustains lifelong olfactory neurogenesis (Caggiano, Kauer, and Hunter 1994; Graziadei and Graziadei 1979; J. E. Schwob et al. 1994). Upon injury, horizontal basal cells (HBCs), the normally quiescent stem cells of the OE, self-renew and differentiate to replace lost neurons, neural progenitors, and other damaged cells, a process requiring the production of multiple cell types in concert in order to repair the damaged system (James E. Schwob, Youngentob, and Mezza 1995; Leung, Coulombe, and Reed 2007; Iwai et al. 2008).

We previously investigated the cellular mechanisms underlying lineage specification during OE homeostasis (Fletcher et al. 2017) and renewal after injury (Gadye et al. 2017) using single-cell RNA sequencing with in vivo lineage tracing of HBCs and their descendants.The molecular pathways that regulate the rapid transition of HBCs from a resting to an activated state have yet to be fully characterized, however. Advances in single-cell profiling have shed light on the role of chromatin accessibility in cell plasticity, cell fate potential, and functional heterogeneity within progenitor cell populations and have improved the ability to predict gene regulatory networks (Trevino et al. 2021; Ma et al. 2020) (reviewed in (Shema, Bernstein, and Buenrostro 2019)). Here, we applied single-cell sequencing to assess the epigenetic and transcriptomic states of individual lineage-traced cells arising from HBCs during recovery from acute injury. The depth and scope of this dataset enabled us to identify transcription factor cascades that are associated with OE regeneration and uncover a subset of HBCs that are poised to respond to injury, as indicated by accessible chromatin around genes that are transcriptionally silent in a subset of quiescent cells but are rapidly expressed shortly after injury. These findings identify a latent activated stem cell state poised to repair the olfactory epithelium in response to injury and contribute to our understanding of the general principles governing neural stem cell maintenance and injury-induced repair of the nervous system.

## Results

### Reconstructed lineages from single cells reveal dynamic transcriptional response to injury

To investigate the transcriptional response to injury at the single-cell level, we analyzed single-cell RNA sequencing data from an injury recovery time course published previously, in which the olfactory epithelium of *Krt5-CreER(T2); Rosa26*^*eYFP/eYFP*^ adult (age 3–7 weeks) mice (23 total; 15 F and 8 M) was injured using methimazole administration, and cells were sampled at 24h, 48h, 96h, 7 days, and 14 days after injury (Gadye et al. 2017; Brann et al. 2020). 20,426 cells were analyzed after filtering (Brann et al. 2020). Uniform Manifold Approximation and Projection (UMAP) dimensionality reduction (McInnes, Healy, and Melville 2018) revealed structure that was correlated with the chronological time of the sampled cells **(Supplementary Figure 1a)**, indicating that the injury triggered a dynamic transcriptional response underlying differentiation. Cell types identified by manual annotation of clusters using expression of known marker genes included activated horizontal basal cells (HBCs*), regenerated HBCs (rHBCs), sustentacular cells (Sus), globose basal cells (GBCs), immature olfactory sensory neurons (iOSNs), and mature olfactory sensory neurons (mOSNs) **(Figure 1a,b)**.

**Figure 1:**
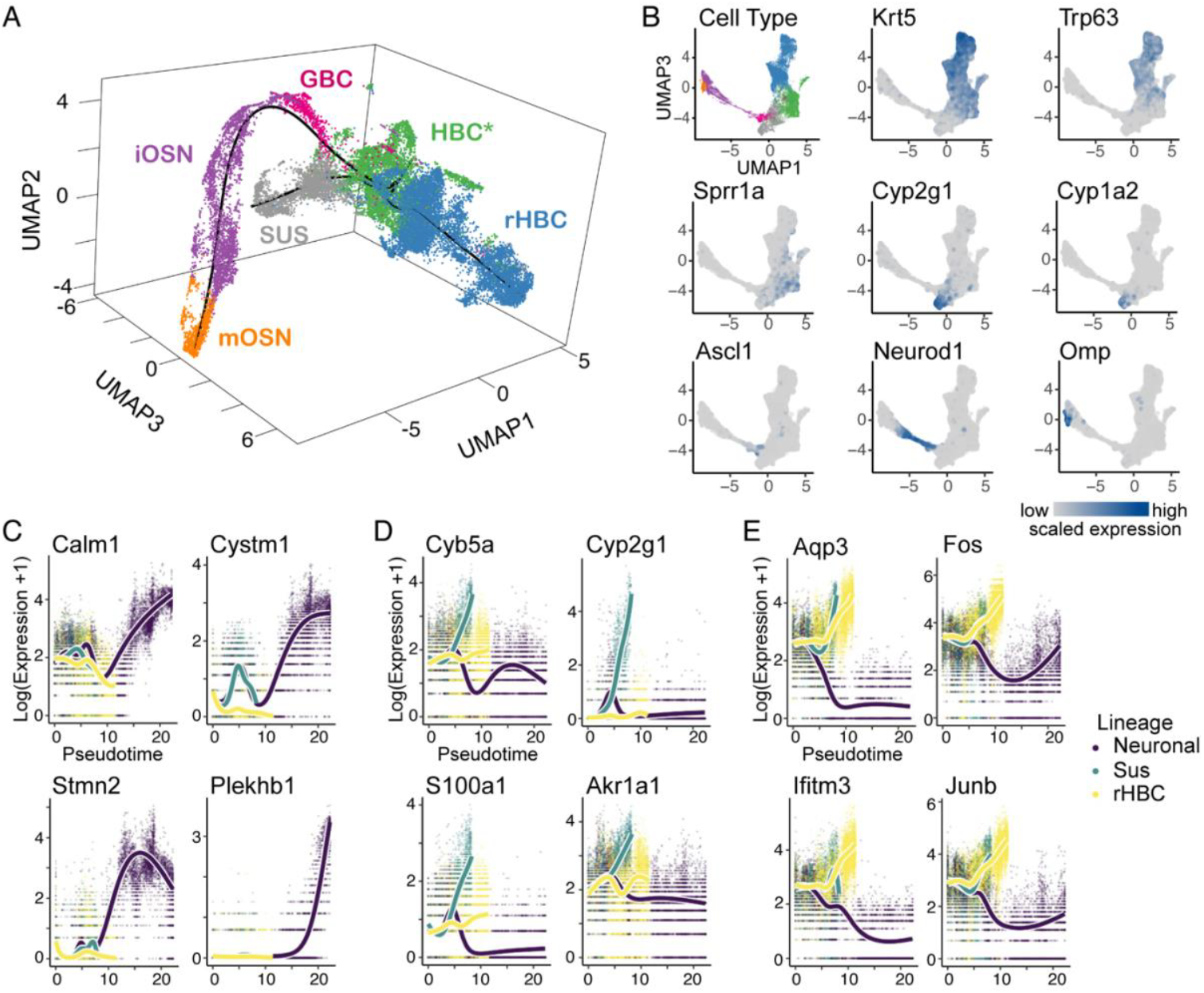
Trajectory inference and differential expression analysis of lineage-traced scRNA-seq data. **(A)** Inferred trajectory in 3D UMAP space, with cells colored according to cell type. Starting from the HBC* population, the trajectory consists of three lineages developing into rHBC, Sus, and mOSN cells. **(B)** Cells in 2D UMAP space, colored according to cell type (top left panel) or the expression of known markers (all other panels), grey denoting no/low expression and blue denoting high expression. **(C-E)** Markers for each lineage identified by differential expression for the neuronal (C), sustentacular (D), and rHBC lineage (E).

To infer the developmental trajectory of HBC progeny, we used *slingshot* (Street et al. 2018) *on UMAP reduced-dimensional space. This analysis revealed a trajectory consisting of three lineages starting from HBCs*: rHBCs, Sus cells, and mOSNs via GBCs and iOSNs (****Figure 1a, black line****). To find genes associated with the development of each cell type, we looked for genes that increased in expression as a function of pseudotime using tradeSeq* trajectory-based differential expression analysis (Van den Berge et al. 2020). This revealed the top genes significantly upregulated in each of the lineages: two of the top four genes in the neuronal lineage are implicated in neuron development and amyotrophic lateral sclerosis (Prudencio et al. 2020; Marques et al. 2020), while three of the top four genes in the Sus lineage are involved in oxidation-reduction, and two of the top four rHBC lineage genes are immediate early genes **(Figure 1c-e, Supplementary Figure 1b-d)**. While all three lineages showed upregulation of the same 1,200 genes following injury, out of a total of 14,618 genes, a particularly large set of genes (5,325) was uniquely involved in the neuronal lineage **(Supplementary Figure 1e)**, possibly reflecting the complex transcriptional programs associated with neurogenesis.

### Employing smooth gene expression profiles to reveal transcription factor expression cascades

The large number of genes activated upon injury suggests that coordinated gene regulatory mechanisms may underlie the transcriptional changes associated with lineage determination. Our previous studies showed gene expression changes specific to Sus and mOSN differentiation to be wave-like or modular, respectively (Fletcher et al. 2017). To further our understanding of the gene regulatory mechanisms underlying differentiation of the three HBC-derived lineages, we sought to identify changes in transcription factor expression along each lineage, focusing on a set of 1,532 mouse transcription factors (TFs) (see Methods). The goal of our approach was to infer which TFs significantly increase in expression along each lineage over the injury recovery time course. Using the fitted gene expression functions along pseudotime from *tradeSeq* (Van den Berge *et al*. 2020), we tested whether the first derivative of the fitted expression function for a given TF was significantly greater than an arbitrary threshold. This allowed us to 1) assess which TFs were significantly peaking at some point along a lineage and 2) derive at which point along differentiation each TF was most active, which here we assume to correspond to the most statistically significant increase in expression.

This analysis uncovered clear TF expression cascades with distinct sequential but gradual activation patterns for each of the lineages **(Figure 2a)**. The number of TFs contributing to each cascade was 524, 231, and 284 for the mOSN, Sus, and rHBC lineages, respectively. We also found 352, 31, and 61 TFs to have a significant peak only in the mOSN (e.g., *Neurog1*), Sus (e.g., *Sec14l2*), or rHBC (e.g., *Trp63*) lineages, respectively **(Supplementary Figure 2b-d; Supplementary File 1**; see Methods). The difference in the number of TFs with a significant peak between the mOSN lineage and the other two lineages may reflect the complexity of the transcriptional programs involved in, and the longer developmental path traversed by, the neuronal lineage. To characterize transcriptional programs along the mOSN lineage, we grouped the 524 TFs in the mOSN cascade according to the moment within the lineage in which they were most active and inferred which cellular processes were activated over time using gene set enrichment analysis. We observed an initial stress response at the early HBC* stage, which was followed by cell cycle regulation and neuron differentiation during the GBC and iOSN stages. A large group of TFs at the iOSN and mOSN stages were involved in processes such as dendrite development, cell projection, and calcium mediated signaling **(Supplementary Figure 2e; top gene ontology terms for each gene set provided in Supplementary File 1)**.

**Figure 2:**
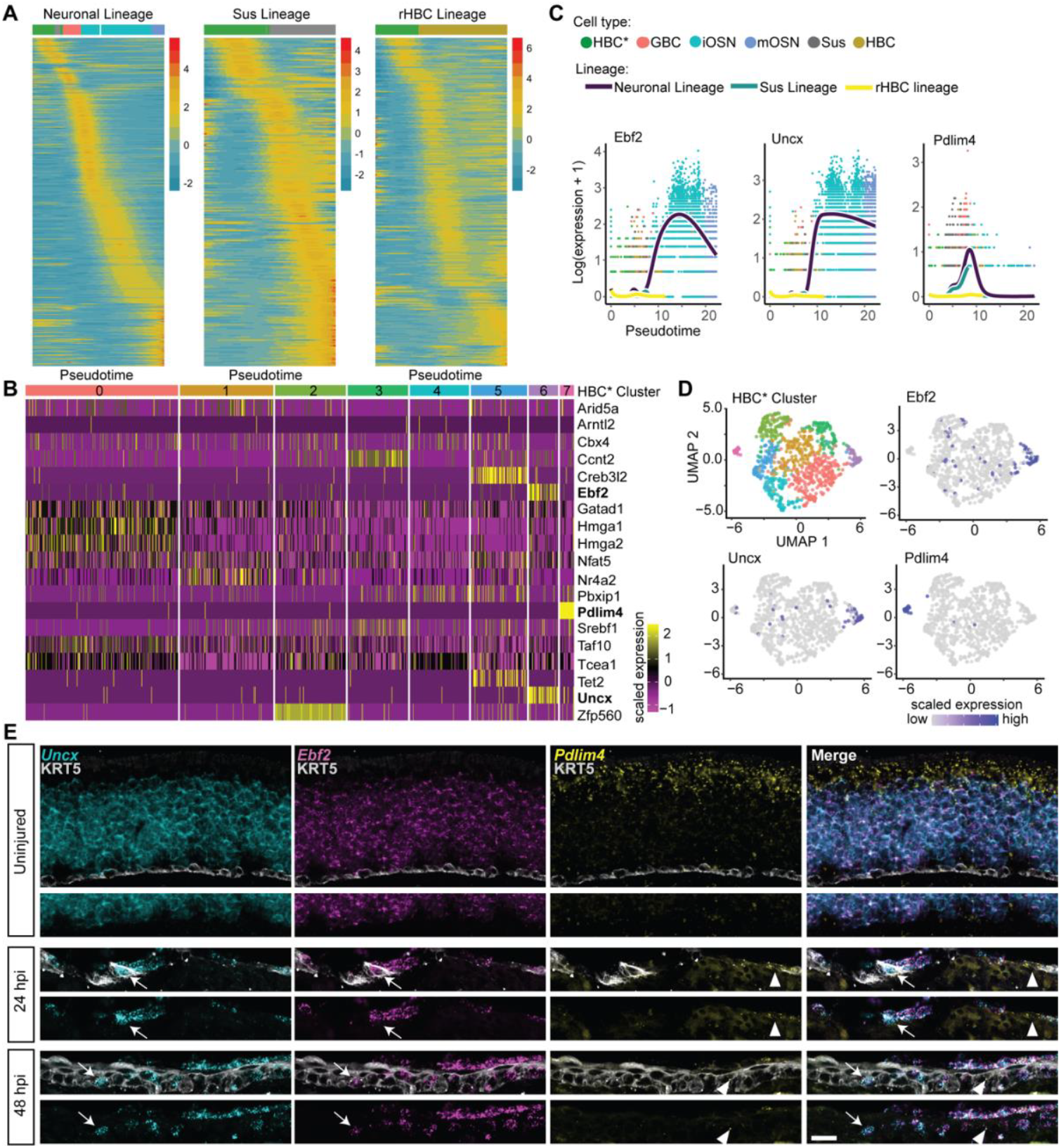
Activated HBCs transiently express lineage-enriched transcription factors. **(A)** Inferred transcription factor (TF) expression cascade for each of the three lineages in the trajectory. Heatmaps of fitted expression measures from tradeSeq, where the x-axis for each panel represents 100 equally wide pseudotime bins for a given lineage, and the most abundant cell type in each bin is indicated using the colorbar at the top of the heatmap (colors correspond to the key in C; If there are too few cells in a bin, no color is provided). Each row in each heatmap represents the expression of a TF normalized to zero mean and unit variance within a lineage. The TFs are ordered according to the pseudotime of their most significant peak, uncovering a TF activity cascade within each lineage. **(B)** Heatmap of the expression of “shared” TFs in activated HBCs clustered on the expression of these 19 TFs. Each row represents the scaled expression of a TF, and the colorbar at the top of the heatmap indicates the cluster label. **(C)** Expression of Ebf2, Uncx, and Pdlim4 (left to right) in each lineage over pseudotime as visualized using tradeSeq. **(D)** Activated HBCs represented in the heatmap in B plotted in 2D UMAP space and colored according to cluster (left) or the expression level of the TF indicated. **(E)** Fluorescent in-situ hybridization (FISH) of Ebf2, Uncx, and Pdlim4 in sections of uninjured, 24 HPI, and 48 HPI OE. Ebf2 is shown in magenta, Uncx in cyan, and Pdlim4 in yellow. Immunohistochemistry using an antibody to KRT5 to mark HBCs is shown in grey.

Previous studies demonstrated that the decision of an HBC to differentiate or self-renew manifests at the stage of HBC activation (Gadye et al. 2017). We therefore hypothesized that differential expression of specific TFs between individual HBCs* could be used to distinguish subpopulations of HBCs* that are committed to different cell fates. To test this hypothesis, we clustered HBCs* sampled at 24 hours post injury (HPI) on the expression of 19 TFs whose expression peaked at the same point in pseudotime during the HBC* stage, prior to the split in lineages **(Supplementary Figure 2a; Supplementary File 1)**. This analysis identified eight discrete clusters **(Figure 2b)**: two clusters were distinguished by specific expression of a single TF (*Zfp560* in cluster 2 and *Pdlim4* in cluster 7) and two clusters by specific co-expression of two TFs (*Creb3l2* and *Tet2* in cluster 5 and *Uncx* and *Ebf2* in cluster 6). These TFs may reflect or even underlie events driving HBCs* toward different cell fates. To test this possibility, we chose three genes whose expression defines two potentially lineage-specific clusters -- cluster 6 (high expression of *Ebf2* and *Uncx*) and cluster 7 (high expression of *Pdlim4*) -- to visualize by tradeSeq and fluorescent in situ hybridization (FISH). We found that *Ebf2* and *Uncx* expression is highest in iOSNs and maintained in mOSNS, and *Pdlim4* expression, while high in GBCs, is maintained only in Sus cells **(Figure 2c)**. FISH confirmed that in the uninjured OE, *Ebf2* and *Uncx* were co-expressed in OSNs and excluded from Sus cells, whereas *Pdlim4* was restricted to Sus cells, consistent with the *tradeSeq* analysis; all three were excluded from resting HBCs **(Figure 2e)**. Consistent with the *cascade* findings, expression of all three genes was evident in putatively activated HBCs by FISH 1-2 days following injury: a subset of HBCs co-expressed *Ebf2* and *Uncx*, whereas a mutually exclusive subset of HBCs expressed *Pdlim4* **(Figure 2e arrows versus arrowheads)**. These findings demonstrate that activated HBCs are transcriptionally heterogeneous as early as 24 HPI and suggest that this heterogeneity could reflect the early adoption of lineage-specific cell fates.

### Estimation of transcription factor activity in activated HBCs through deconvolution of mRNA expression

To further distinguish which TFs might be involved in activating gene expression during olfactory neurogenesis, we started by identifying the genes that are expressed along the mOSN lineage. We then inferred which of the 524 TFs in the mOSN cascade (shown in **Figure 2a**) were likely to drive their expression (see Methods). Briefly, we resolved the collective change in gene expression over pseudotime along the lineage into co-regulated gene groups. Next, for each cell in the lineage and each of the 524 mOSN cascade TFs, we probabilistically assigned mRNAs to the TFs that most likely drove their expression. Finally, we estimated TF activity based on the number of mRNAs likely driven by each TF within bins of pseudotime. This revealed 3 clusters of TFs that were most active either at the HBC* (early), GBC (mid), or iOSN-mOSN (late) stage **(Figure 3a)**. A closer look at the 20 most variable TFs in terms of their activity measure **(Figure 3b)** revealed that several known differentiation genes, including *Sox11* and *Rfx3*, were most active late in the lineage, while *Hes6, Ezh2*, and *E2f1* were most active at the GBC stage. Additionally, the immediate early genes *Egr1, Fos, Junb*, and *Jund* and the pioneer TF *Foxa1* were most active early in the lineage **(Figure 3c)**. Using multiplex FISH, we confirmed increased expression of *E2f1, Ezh2, Sox11*, and *Rfx3* in the regenerating OE at 96 HPI, just after the split in the OSN and Sus lineages **(Figure 3d)**. In addition, we confirmed increased *Egr1* and *Fos*, which were estimated to be active at the HBC* stage, in activated HBCs, identified by high *Lgals1* or high *Krt5* expression, at 48 HPI **(Figure 3e)**. Finally, we confirmed increased expression of *Foxa1*, and lack of expression of the homologous *Foxa2* and *Foxa3*, at 48 HPI **(Figure 3e)**.

**Figure 3:**
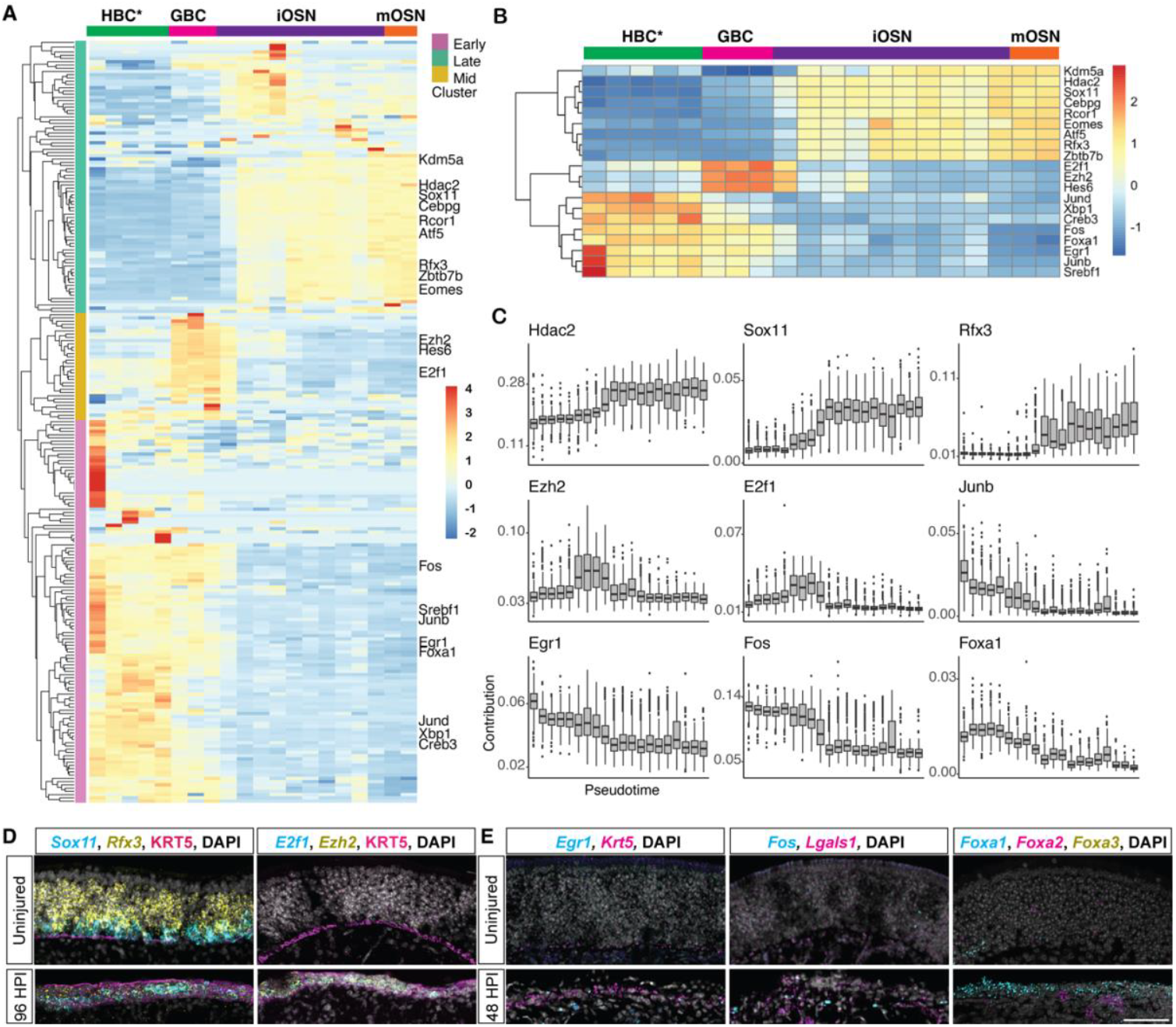
Deconvolution of gene expression reveals dynamic TF activity along the mOSN lineage. **(A)** Clustered heatmap of scaled activity for TFs (rows) that are differentially active along the mOSN lineage. The x-axis represents pseudotime and the dominant cell type in each pseudotime bin is indicated at the top of the heatmap. Hierarchical clustering revealed three groups of TFs: early (mauve), mid (amber), and late (green). **(B)** Clustered heatmap of scaled TF activity for the 20 most variable TFs. **(C)** Change in TF activity over pseudotime for representative TFs from each cluster. **(D)** Fluorescent in situ hybridization (FISH) of Sox11 (cyan), and Rfx3 (yellow) or E2f1 (cyan) and Ezh2 (yellow) showing increased expression in regenerating OE at 96 HPI. HBCs were identified with an antibody to KRT5 (magenta) and nuclei with DAPI (grey). **(E)** FISH of two TFs active at the HBC* stage, Egr1 and Fos (cyan), showing increased expression in activated HBCs (high Krt5 or Lgals1, magenta) at 48 HPI compared with before injury, and FISH of Foxa1 (cyan), Foxa2 (magenta), and Foxa3 (yellow) showing increased expression of Foxa1 in basal cells at 48 HPI (magenta, yellow signal is non-specific). Nuclei labeled with DAPI (grey). Scale bar in E for D,E = 50 microns

### Epigenetic priming of mRNA transcription in the HBC injury response

The rapid and coordinated up-regulation of gene expression in HBCs in response to injury led us to hypothesize that HBCs are epigenetically primed to initiate rapid injury-induced changes in mRNA transcription, as observed in other stem cell niches, such as muscle stem cells (Rodgers et al. 2014), resting CD4+ T cells (Z. Wang et al. 2009; Rogers et al. 2021), hepatocytes (Reizel et al. 2021), and Müller glia (Norrie et al. 2024). To test this hypothesis, and to identify gene regulatory networks that are initiated upon injury, we performed an assay for transposase accessible chromatin with sequencing (ATAC-seq) (Buenrostro et al. 2013; Corces et al. 2017) on ICAM1+ HBCs isolated either from uninjured olfactory epithelium (hereafter referred to as “uninjured HBCs”) or from olfactory epithelium 24 hours after injury with methimazole (hereafter referred to as “injured HBCs”). We also generated bulk RNA sequencing libraries from uninjured and injured HBCs isolated from additional samples to serve as a gene expression reference.

As a control, we first asked whether gene expression was correlated with chromatin accessibility in uninjured HBCs. To this end, we established two gene sets: 1) genes that are highly expressed in HBCs and 2) olfactory receptor genes, which are not expressed (“silent”) in uninjured HBCs. We then assessed chromatin accessibility in the regions surrounding these genes’ transcriptional start sites (TSS) in uninjured HBCs **(Figure 4a)**. As expected, regions surrounding the TSS of highly expressed genes were found in accessible chromatin, whereas those of silent olfactory receptor genes did not appear in accessible chromatin, demonstrating that the TSS of these silent genes generally occupies inaccessible chromatin.

**Figure 4.**
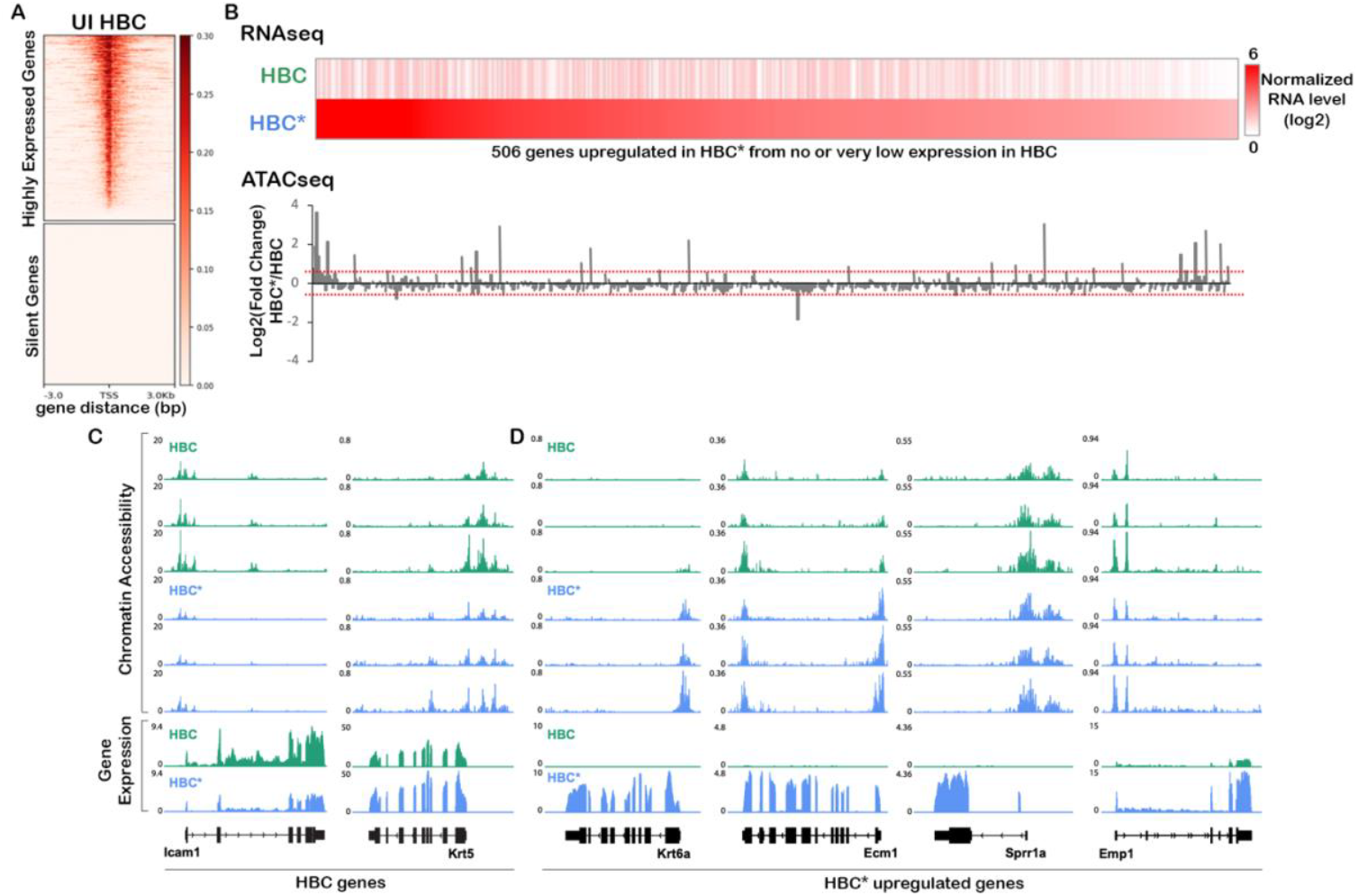
Early response genes are primed for activation at the chromatin level. **(A)** Heatmaps of ATAC-seq read counts from uninjured (UI) HBCs around the TSS of highly expressed genes (top) or silent olfactory receptor genes (bottom). **(B)** Heatmap of normalized bulk gene expression values (average RPKM of two biological replicates) in uninjured HBCs versus injured HBCs (HBC*) for the top 506 genes that are upregulated in HBCs after injury, with genes ordered from left to right according to descending expression in injured HBCs (top). Bar graph showing the log2(fold-change) in chromatin accessibility after injury relative to before injury (bottom), where dotted lines indicate log2(fold-change) of 0.5 and -0.5. **(C,D)** ATAC-seq (top) and bulk RNA-seq (bottom) read counts before (green) and after (blue) injury around genes that are known to decrease (Icam1) or increase (Krt5) in expression in injured HBCs (C) and wound response genes (Krt6a, Ecm1, Sprr1a, and Emp1) (D).

We then identified epigenetic changes associated with injury. Bulk RNA-seq data were used to define 506 “early response” genes that were upregulated in injured HBCs and had no or very low expression in uninjured HBCs. We then quantified changes in chromatin accessibility around each early response gene’s TSS between uninjured and injured HBCs. Interestingly, we found that these early response genes tended to have comparable accessibility around the TSS both before and after injury despite significant increases in gene expression, suggesting that some of these genes may be primed for rapid activation upon injury **(Figure 4b)**. We next looked at chromatin changes after injury over the gene bodies of individual marker genes for HBCs (*Icam1* and *Krt5*) and wound response genes identified and defined in (Gadye et al. 2017; Brann et al. 2020)) (*Krt6a, Ecm1, Sprr1a*, and *Emp1*). We found that *Icam1* lost and *Krt5* gained chromatin accessibility consistent with changes in gene expression upon injury. Similarly, the chromatin around the wound response gene *Krt6a* became accessible exclusively when *Krt6a* transcription was activated. However, other wound response genes already exhibited partially (e.g., *Ecm1*) or fully (e.g., *Sprr1a* and *Emp1*) accessible chromatin prior to injury, suggesting that they may be primed for activation upon injury **(Figure 4c,d)**.

### Single-cell analysis of chromatin accessibility reveals multiple HBC states

We next asked whether wound response genes are epigenetically primed in all or a subset of uninjured HBCs by performing single-cell ATAC-seq on lineage-traced FACS purified HBCs (*Krt5-CreER(T2)*; *Rosa26*^*eYFP/eYFP*^) before and 24 hours after injury with methimazole. Chromatin accessibility profiles of 5,743 cells were obtained. Gene activity scores were estimated using the ArchR framework for scATAC-seq (Granja et al. 2021). After quality control and doublet removal, 4,732 cells remained. Batch-corrected Harmony embeddings (Korsunsky et al. 2019) were used as input to UMAP dimensionality reduction for visualization. Clustering on the Harmony embeddings identified the major cell types that were present in the scRNA-seq data. HBCs made up the largest cluster; neuronal and sustentacular cells were identified by the open chromatin state of their respective marker genes **(Supplementary Figure 3a,b)** and removed from subsequent analysis. Interestingly, the remaining subset of 4,076 HBCs consisted of three distinct clusters, possibly corresponding to different HBC states **(Figure 5a)**: two clusters consisted almost exclusively of either injured or uninjured cells, which corresponded to activated and resting (or quiescent) states, respectively, as defined previously by scRNA-seq (Gadye et al. 2017; Fletcher et al. 2017). However, a third cluster contained nearly equal numbers of injured and uninjured cells **(Figure 5b,c)**, which may reflect a distinct epigenetic state shared by a subset of both resting and activated HBCs. Given the mixture of injured and uninjured cells, we labeled this third cluster “hybrid.” To distinguish between the experimental origin of HBCs and the predicted HBC state going forward, we use the terms “injured” and “uninjured” when referring to experimental origin, and “resting,” “activated,” and “hybrid” when referring to inferred cell state.

**Figure 5:**
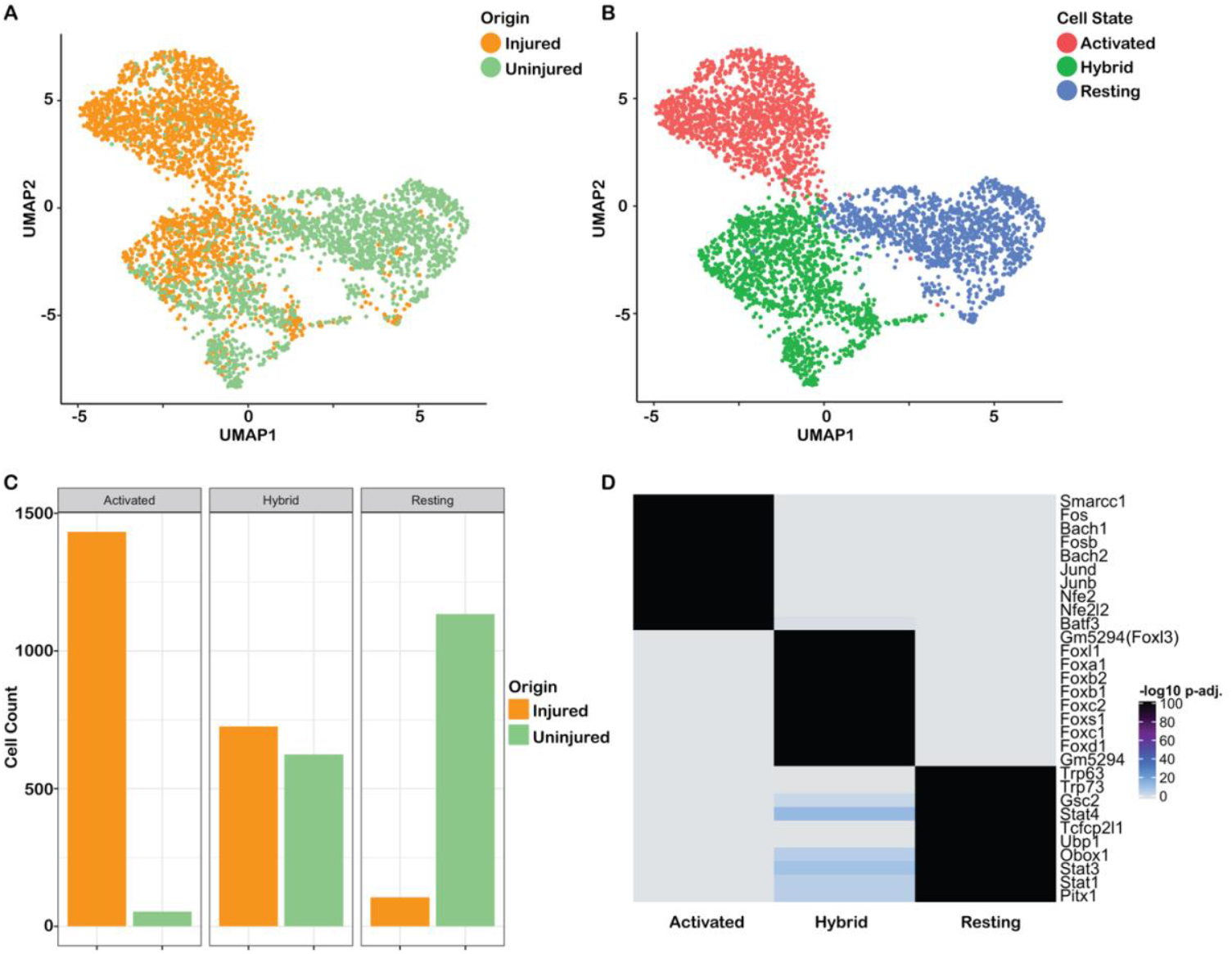
scATAC-seq data uncover three HBC states. **(A-B)** UMAP dimensionality reduction of the scATAC-seq data; cells are colored according to origin (A) or according to state (B). **(C)** Barplot visualizing the number of (un)injured cells in each cluster. **(D)** Heatmap (unclustered) showing the Benjamini-Hochberg FDR adjusted p-values for testing the null hypothesis of no enrichment, of the top 10 enriched TF motifs (rows) for each cell state (columns). The heatmap clearly illustrates that the three sets of TFs are high/insignificant (grey) in specific HBC groups and low/significant (black) in others.

To assess in which state(s) wound response genes are accessible, we visualized the normalized accessibility of two HBC marker genes (*Icam1* and *Krt5*) and the four wound response genes assessed above (*Krt6a, Ecm1, Sprr1a*, and *Emp1*) in the three scATAC-seq clusters by UMAP and density plot **(Supplementary Figure 4)**. *Icam1* was most accessible in the resting HBC state and least accessible in the activated HBC state, consistent with its downregulation upon injury **(Supplementary Figure 4a-c,e)**. Similarly, *Krt5* was most accessible in the activated HBC state, consistent with its upregulated expression in HBCs upon injury **(Supplementary Figure 4a,d,f)**. Among the wound response genes, *Krt6a* and *Ecm1* were most accessible in the activated state, and more accessible in the hybrid state compared with the resting state. *Sprr1a* and *Emp1*, which the bulk ATAC-seq data predicted to be accessible in uninjured HBCs, were approximately equally more accessible in both the activated and hybrid states than the resting state **(Supplementary Figure 4a,d,f)**, suggesting that a subset of HBCs from uninjured OE (the “hybrid” HBCs) are poised to activate a subset of wound response genes prior to injury.

To determine whether the three HBC clusters indeed represent distinct epigenetic states, we looked for regions of accessible chromatin that distinguished each cluster. Peak calling using MACS2 (Zhang et al. 2008) on the three HBC clusters resulted in a set of 183,712 peaks. We identified marker peaks for each of these clusters using *ArchR* (see Methods), uncovering 12,146, 8,492, and 3,513 marker peaks for the activated, hybrid, and resting clusters, respectively (nominal FDR <= 0.01 and log_2_FC > 1.25). The cisbp database (Weirauch et al. 2014) was used to derive TF binding site motifs, which were subsequently assessed for enrichment in these sets of marker peaks **(Figure 5d)**. The results for the activated and resting clusters reflected TFs hypothesized to play a role in HBC activation or quiescence, respectively: motifs for the activity-responsive TFs *Fosb* and *Junb* were enriched in the activated state, and motifs for the canonical resting HBC TFs *Trp63* and *Trp73* were enriched in the resting state. Also enriched in activated HBCs were motifs for *Smarcc1*, the main regulatory component of the Brg1-associated factor (BAF) chromatin remodeling complex important for early stem cell differentiation, cell fate determination, and keratin gene expression (Lim et al. 2020; Schaniel et al. 2009). Remarkably, the top upregulated motifs for the hybrid cell state were all members of the Forkhead box (Fox) family of TFs.

### Integration of scRNA-seq and scATAC-seq data suggests a latent activated olfactory stem cell state

In light of our discovery of multiple HBC epigenetic states, we were interested in linking these states as revealed by scATAC-seq with the transcriptomic states identified by scRNA-seq. Our two corresponding scRNA-seq sources for the two scATAC-seq HBC sources (injured and uninjured) were: 1) cells sampled at 24 HPI and clustered at the starting point of the trajectory (i.e., contained within the cluster that was assigned as the starting cluster) and 2) regenerated HBCs taken from the 14 days post-injury timepoint, which are indistinguishable from uninjured HBCs at the transcriptional level (Brann et al. 2020; GEO accession GSE173999) **(Supplementary Figure 5a,b)**. When we examined these HBC transcriptomes by UMAP, we found that regenerated/resting HBCs formed a single cluster and HBCs at 24 HPI formed three clusters (a primary cluster and two smaller clusters) **(Figure 6a)**. Next, to link epigenetic and transcriptomic states, we identified gene activity markers for each of the scATAC-seq clusters and mapped them onto the scRNA-seq clusters. Importantly, we first confirmed that both injured and uninjured cells contributed to the gene activity markers of the hybrid cell state **(Supplementary Figure 5b)**. We then visualized the gene activity markers for each HBC state in the scRNA-seq data, which revealed enrichment of marker gene expression in the analogous scRNA-seq cell clusters (activated markers in the main 24 HPI cluster and resting markers in the regenerated cluster) **(Figure 6b,d)**, linking these cells in epigenetic and transcriptomic space. Interestingly, the hybrid marker genes were enriched in the two smaller 24 HPI clusters, suggesting that hybrid genes, while accessible in both injured and uninjured cells, are activated upon injury in a small subset of HBCs **(Figure 6c)**.

**Figure 6:**
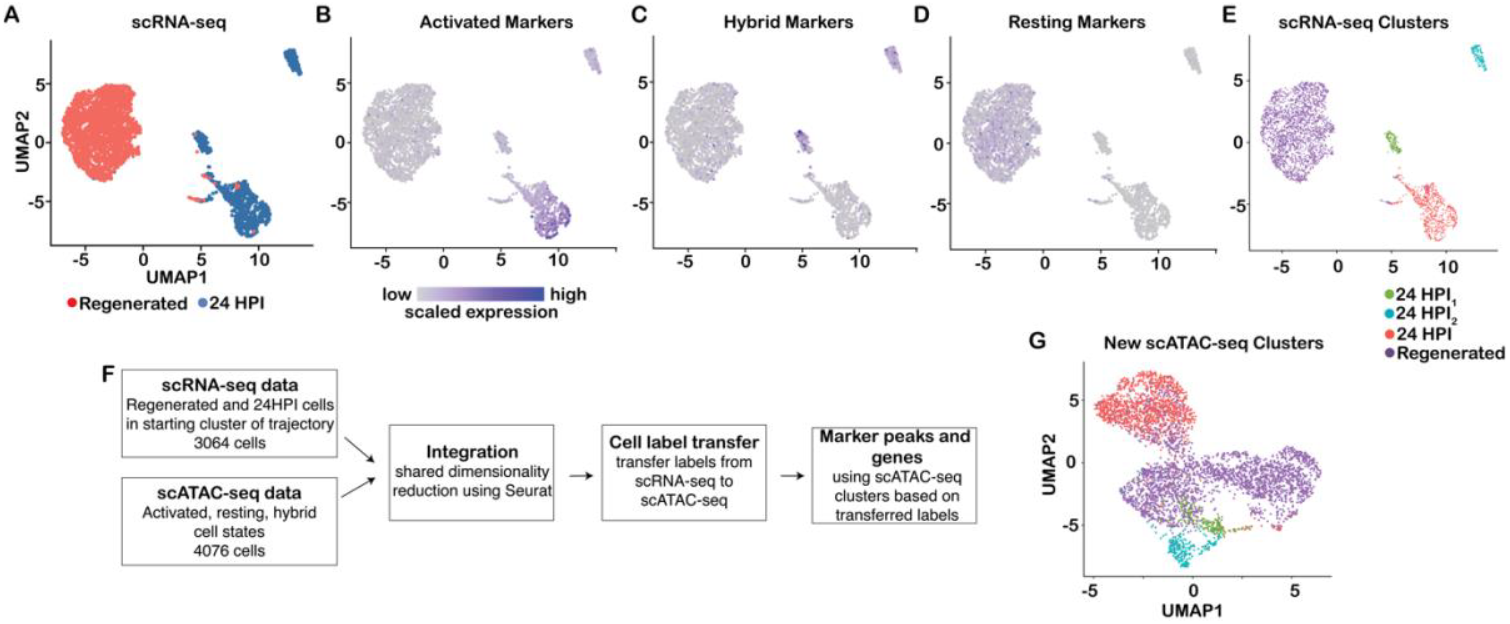
Integration of scRNA-seq and scATAC-seq data. **(A-D)** scRNA-seq data of activated and regenerated HBCs visualized in UMAP space. (A) scRNA-seq data with cells colored according to cell type. (B-D) scRNA-seq data with cells colored according to expression of genes that were found to be markers for each of the three cell states identified using scATAC-seq gene activity scores. ‘Activated’ and ‘Resting’ genes identified using scATAC-seq are correspondingly upregulated in scRNA-seq data for activated cells (B) and regenerated cells (D), while ‘Hybrid’ genes are upregulated in the two small activated subclusters (C). The expression values were first scaled within each gene to have zero mean and unit variance across all cells, upon which the scaled expression was summed across genes within each cell. **(E)** scRNA-seq data with cells colored according to cell state. **(F)** Workflow for cell label transfer from scRNA-seq to scATAC-seq data. First, Seurat was applied to integrate the scRNA- seq and scATAC-seq data by shared dimensionality reduction using canonical correlation and to transfer the scRNA-seq cell labels to the scATAC-seq dataset. Next, the transferred labels are used to define marker peaks and genes in the scATAC-seq dataset. **(G)** scATAC-seq data with cells colored according to cell state predicted by transferring cell labels from scRNA-seq data.

Given this link between accessible chromatin and gene expression, we used the Seurat workflow (Stuart et al. 2019) to integrate the scRNA-seq and scATAC-seq datasets. After shared dimensionality reduction **(Supplementary Figure 5c)**, we applied cell type label transfer analysis, where manual cell type annotations based on the scRNA-seq (*regenerated, 24 HPI, 24 HPI*_*1*_, and *24 HPI*_*2*_) clusters were transferred to scATAC-seq data, to predict a corresponding cell type in the latter. This revealed that the *24 HPI*_*1*_ and *24 HPI*_*2*_ sub-clusters were recovered within the scATAC-seq hybrid cluster, and they were tightly compacted within the reduced-dimensional space **(Figure 6 e-g)**.

We then searched for marker genes and TF motifs for each of the four new scATAC-seq clusters identified after label transfer using *ArchR* (see Methods) **(Supplementary Figure 6a,b)**. Marker genes appeared similar between the *24 HPI*_*1*_ and *24 HPI*_*2*_ clusters, while the *24 HPI* and *regenerated/resting* clusters were readily distinguishable. Moreover, the top enriched TF motifs were similar between *24 HPI*_*1*_ and *24 HPI*_*2*_ and corresponded to the motifs identified previously for the main hybrid state (shown in **Figure 5d**), suggesting that *24 HPI*_*1*_ and *24 HPI*_*2*_ are subclusters of the main hybrid cluster. The enrichment results again pointed to a heavy involvement of Fox TFs for the two hybrid subclusters, which was supported by higher expression of the relevant Fox TFs (*Foxa1, Foxb1, Foxb2, Foxc1, Foxc2, Foxd1, Foxl1*, and *Foxs1*) in the *24 HPI*_*1*_ and *24 HPI*_*2*_ scRNA-seq clusters **(Supplementary Figure 6c,d)**.

### Hybrid-specific ATAC-seq peaks are associated with genes expressed in HBCs only after injury

Transcription of lineage-specific genes has been shown to be primed in quiescent progenitor cells in several adult stem cell niches, including liver (Reizel et al. 2021; Iwafuchi-Doi et al. 2016), bone marrow (Paul et al. 2015), skeletal muscle (Okafor et al. 2023), and naive CD4+ T cells (Rogers et al. 2021). Moreover, transcriptional priming is mediated by Foxa TFs in multiple endoderm-derived tissues (A. Wang et al. 2015; Iwafuchi-Doi et al. 2016; K. Lee et al. 2019; Geusz et al. 2021). Given the early expression of the pioneer TF *Foxa1* in the neuronal lineage and the enrichment of Fox motifs in hybrid-specific ATAC-seq peaks, we investigated whether hybrid HBCs have accessible chromatin around lineage-specific genes. We ranked the marker genes for each ATAC-seq subcluster (identified using ArchR above) by FDR **(Supplementary File 2)** and visualized high-ranking known lineage-specific genes by UMAP and density plot. We found that the activity scores for known markers of Sus (*Cd36, Aldh1a7*), OSNs (*Trim46, Erich3, Cap2*), and regenerated HBCs (*Adh7*) were enriched in the hybrid cluster **(Supplementary Figure 7a)** and confirmed that the distribution of activity scores for each of these genes was approximately equal for injured and uninjured hybrid cells **(Supplementary Figure 7b, Supplementary File 2)**. Moreover, we visualized lineage-enriched expression of each of these genes using *tradeSeq* **(Figure Supplementary Figure 7c)**. These results reveal that a subset of silent lineage-specific genes occupy open chromatin in some uninjured HBCs.

We were also interested in whether distal ATAC-seq peaks specific to the hybrid cluster were associated with genes whose priming would facilitate lineage specification and/or epithelial regeneration. While ArchR prioritizes local accessibility, Genome Region Enrichment of Annotations Tool (GREAT) assigns peaks to genes without assigning weight based on proximity to the gene body and searches for enriched biological functions (McLean et al. 2010). To expand our search for hybrid genes to include distal peak-gene associations, we used GREAT with default parameters. GREAT associated the 8,492 differentially accessible hybrid peaks, the majority of which were distal (>50 kb) to the transcription start site, with 6,426 genes. Of these, 424 were significantly enriched for hybrid peaks (Binomial test, FDR *q-value* < 0.05), including five of the six genes identified by ArchR and reported above (*Adh7, Aldh1a7, Cap2, Cd36*, and *Trim46*). Among the top 100 genes, we identified several candidates with reported roles in epithelial regeneration -- *Foxa1, Klf5, Ets2, Ehf*, and *Sgk1* (full rankings and statistics available in **Supplementary file 2**) -- which we validated by FISH (Paranjapye et al. 2020; Bell et al. 2013; Naik et al. 2017; Ge et al. 2017; Fossum et al. 2017, 2014; Stephens et al. 2013). We found that *Klf5, Ets2, Ehf*, and *Sgk1* transcripts were absent in uninjured HBCs (**Figure 7c**, HBCs identified by Krt5 IHC in blue), but visible in the OSN and Sus layers, which was consistent with our scRNA-seq data (**Figure 7a**). Moreover, all five transcripts were upregulated broadly in HBCs 24 and 48 hours after injury (**Figure 7c**). In addition, despite their notable absence in uninjured HBCs and broad upregulation after injury, all five of these genes were most accessible in the hybrid scATAC-seq cluster, as demonstrated by the median of the distribution of their gene activity scores in the resting, hybrid, and activated scATAC-seq clusters (**Figure 7b**).

**Figure 7:**
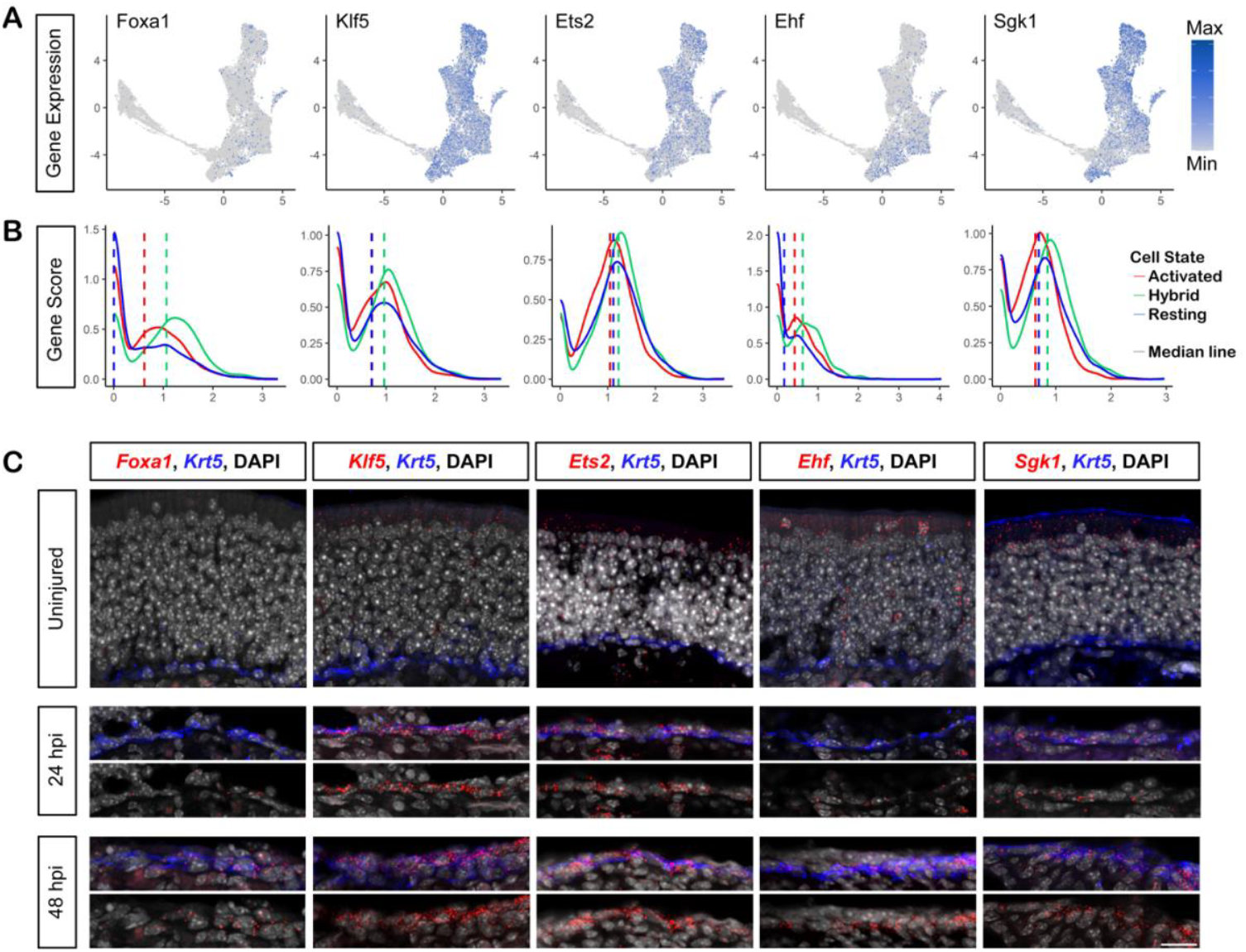
Genes proximal to regions of differentially accessible chromatin in hybrid cells are upregulated following injury. **(A)** Cells in 2D UMAP space, colored according to the expression of selected genes proximal to hybrid cell differentially accessible chromatin, gray denoting no/low expression and blue denoting high expression. The cells at the upper right of the UMAP represent the rHBC lineage, the extended arc of cells toward the left the neuronal lineage, and the cells at the bottom the sustentacular lineage. **(B)** Density plots of normalized gene activity scores for lineage-specific genes in resting (blue), hybrid (green), and activated HBCs. **(C)** Fluorescent in-situ hybridization (FISH) for each gene in (top to bottom) uninjured OE, 24 hours post injury and 48 hours post injury. FISH for each gene shown in red, Krt5 immunohistochemistry shown in blue, and DAPI in grey. Scale bar = 20µM.

Sets of genes associated with differentially accessible hybrid peaks appeared to be enriched for epithelial regenerative functions. Top GO biological processes identified using GREAT included the terms *tissue remodeling* and *lung epithelium development* (**Supplementary file 2**). The most enriched molecular function was cytokine binding, which facilitates epithelial regeneration in OE and other systems (Ullah, Rowan, and Lane 2024; Guenin-Mace, Konieczny, and Naik 2023). By comparison, the top term for resting peaks was *stem cell population maintenance* and for activated peaks was *cell junction organization*. Together, these observations are consistent with the hypothesis that hybrid HBCs are poised for activation through accessible chromatin around injury response and lineage-associated genes that are rapidly upregulated early in the regeneration process.

## Discussion

An adult stem cell’s epigenetic landscape contributes to its ability to respond to injury and regenerate discrete lineages. In the case of the olfactory epithelium stem cell niche, the specific gene regulatory pathways involved in triggering olfactory stem cell activation and determining cell fate are still unclear. In the present study, we used single-cell transcriptomic and epigenetic profiling of HBCs and their descendants to determine the transcription factors that may contribute to cell fate specification and differentiation after injury. We identified TF binding motifs in accessible chromatin that reflect known and novel gene regulatory networks for resting and activated stem cell states. In the process, we further identified a subset of quiescent HBCs with an epigenetic profile similar to a subset of injury-activated HBCs. Collectively called “hybrid” cells, these HBCs represent a novel stem cell state characterized by open chromatin over a subset of wound response and regeneration genes that are only expressed later in the regeneration process.

### Transcriptional cascades define differentiation of multiple lineages in the regenerating olfactory epithelium

We previously reported that HBCs expressing the TF *Hopx* 24 hours after injury are restricted to the mOSN or Sus lineages (but not the regenerating HBC lineage), suggesting that cell fate is chosen in this early heterogeneous activated state (Gadye et al. 2017). Here, we identified 19 TFs that peak in expression in the activated state, some of which are mOSN or Sus lineage markers and define specific clusters or subsets of activated HBCs. In support of this computational analysis, transcripts for the mOSN or Sus marker genes *Ebf2*+*Uncx* or *Pdlim4* were detected by FISH in mutually exclusive activated HBCs 24-48 hours after injury but were not detected in uninjured HBCs.

We hypothesize that this initial transcriptional heterogeneity in gene expression resolves activated HBCs into lineage-committed cells. Accordingly, our prediction of TF cascades should identify the TFs involved in each lineage and the point at which they peak -- and therefore act -- along the lineage. For the mOSN lineage in particular, this function is useful for understanding which TFs are important for regulating specific aspects of neuronal differentiation, from the initial HBC* stage through the final mOSN stage. Thus, we further predicted which TFs are *most* active at each stage by grouping the expression of putative target genes into co-regulated sets, and then inferring which TFs were most likely to drive the expression of each set. This approach identified the TFs that are likely to be the most active at the HBC*, GBC, or OSN stage. Among the putative TFs most active at the HBC* stage, FoxA1 is notable for its role as a pioneer TF in other epithelia (reviewed in (Iwafuchi-Doi and Zaret 2016)) and directly links the FoxA motifs found to be enriched in open chromatin specific to hybrid HBCs with gene expression in activated HBCs.

### Epigenetic heterogeneity of quiescent olfactory stem cells

Recent studies have identified heterogeneity in various adult stem cell populations, including skeletal muscle stem cells (Der Vartanian et al. 2019; Scaramozza et al. 2019; Okafor et al. 2023), thymus epithelial progenitor cells (Rogers et al. 2021; Kadouri et al. 2020; Nusser et al. 2022), hair follicle stem cells (Ma et al. 2020; Yang et al. 2017; Joost et al. 2020), and hematopoietic stem cells (Paul et al. 2015; Perié et al. 2015; Rodriguez-Fraticelli et al. 2018). The hybrid HBC state adds an epigenetic layer of heterogeneity to HBCs. Advances in detecting such heterogeneity owe in part to advances in single-cell epigenetic profiling. Indeed, it has been suggested that early in differentiation, chromatin state may be more effective than gene expression at predicting a progenitor cell’s fate (Ma et al. 2020). It therefore stands to reason that chromatin state may be a particularly good predictor of differences between seemingly homogeneous cells, and may explain why uninjured (quiescent) and injured (activated) HBC populations together were observed to occupy two discrete states transcriptionally, while a third hybrid state was only detected at the epigenetic level.

### A latent activated state may prime olfactory stem cells for regeneration

Our discovery of a hybrid HBC population common to both quiescent and activated cells, which exhibits accessible chromatin around a subset of wound response genes like *Emp1* and *Sprr1a*, suggests that a subset of quiescent HBCs exists in a latent activated state, poised for activation in response to injury. The enrichment of FoxA TF family motifs in hybrid ATAC-seq peaks is notable given that FoxA TFs are associated with enhancer priming (Geusz et al. 2021) and FoxA TFs are established pioneer transcription factors in the regeneration of other epithelia (reviewed in (Iwafuchi-Doi and Zaret 2016)). The strong enrichment of FoxA motifs in accessible chromatin specific to hybrid HBCs could indicate epigenetic priming at a subset of hybrid-specific *cis*-regulatory elements in a manner analogous to the priming of genes in embryonic stem cells (ESCs) by the histone methylation function of Polycomb Repressive Complex 2 (PRC2).

Indeed, throughout the genome of ESCs, regions that harbor both active and repressive histone marks, termed “bivalent domains,” are enriched for differentiation genes, many of which exhibit promoter co-occupancy by PRC2 and the ESC pioneer TFs *Oct4, Sox2*, and *Nanog*, keeping primed differentiation genes silent but poised for transcriptional activation (T. I. Lee et al. 2006; Azuara et al. 2006) (Bernstein et al. 2006). Additionally, RNA polymerase II (RNAPII) stalling at promoters of highly regulated genes provides a mechanism for maintaining a gene’s potential to be reactivated (reviewed in (Core and Adelman 2019)). It should be noted, however, that in mouse ESCs, promoter proximal pausing of RNAPII was found to be enriched at cell cycle and signal transduction genes, rather than differentiation genes, and pausing was found to be required for responsiveness to differentiation cues (Williams et al. 2015). A similar combination of FoxA1-mediated epigenetic priming of differentiation genes and RNAPII stalling at promoters of proliferation genes in HBCs could keep these genes silent but poised for activation upon injury.

In the present study, we identified a discrete population or state of HBCs that may be similarly poised -- perhaps via pioneer TFs at a specific subset of genes -- for later activation of specific genes upon injury. This latent activated state may be a means of maintaining the ongoing balance of quiescence versus proliferation in olfactory epithelium stem cells. A “hybrid” state that is poised for activation in response to injury may therefore provide a strategy for allowing a robust regenerative response in the niche while maintaining stem cell quiescence under homeostatic conditions.

## Acknowledgments

The authors are grateful for ongoing discussions with members of the Ngai, Purdom, and Dudoit groups, particularly Dr. Jonathan Lovas and Hao Wang. This work was supported by the Division of Intramural Research of the National Institute of Neurological Disorders and Stroke, National Institutes of Health (ZIA NS009423) (J.N.) and grant R01 DC007235 from the National Institute on Deafness and Other Communication Disorders, National Institutes of Health (S.D.). K.V.d.B. also acknowledges funding from the Research Foundation Flanders grants 1246220N and V411821N, and the Belgian American Educational Foundation. The content is solely the responsibility of the authors and does not necessarily represent the official views of the National Institutes of Health.

## Methods

### Animals

All animal work was carried out in compliance with the University of California Institutional Animal Care and Use Committee (IACUC) and National Institute of Neurological Disorders and Stroke ACUC according to federal guidelines. Mice containing the *Krt5-CreER(T2)* driver (Indra et al. 1999) and *Rosa26*^*eYFP*^ reporter (Srinivas et al. 2001) were kept on a mixed C57Bl/6J and 129 background. Wild-type *C57BL/6J* mice were purchased from the Jackson Laboratory. Both male and female mice were used in all studies.

### scRNA-seq with lineage tracing

#### OE injury, dissociation, and Fluorescence Activated Cell Sorting (FACS)

HBCs were labeled and their descendents post-injury were lineage traced using the *Krt5- CreER* driver crossed with the *Rosa26*^*eYFP*^ fluorescent reporter, as described previously ((Fletcher et al. 2017); (Gadye et al. 2017)). Briefly, *Krt5-CreER*; *Rosa26*^*eYFP/eYFP*^ mice (23 total; age 3–7 weeks; 15 F and 8 M) were injected intraperitoneally once with tamoxifen (0.25 mg tamoxifen/g body weight) after weaning, then injected with methimazole (50 μg/g body weight, IP) at least one day after tamoxifen administration, and sacrificed at 24 h, 48 h, 96 h, 7 d, or 14 d after injury with methimazole. For each experimental time point, the dorsal olfactory epithelium (OE) was surgically removed and dissociated, as described in Fletcher et al. 2017 and Gadye et al. 2017: OE from each animal was individually processed in approximately 1 mL of pre-warmed (37°C) dissociation medium (150 units papain dissolved in 5 mL Neurobasal medium with 2.5 mM Cysteine and 2.5 mM ethylenediaminetetraacetic acid) with 100 units DNAse I and incubated at 37°C for 25 mins. Samples were then washed three times with 10% fetal bovine serum in phosphate buffered saline (PBS-FBS) and strained through a 35 μm nylon mesh filter cap into a 5 mL polypropylene tube to remove debris. Propidium iodide (PI) was added to cells at a final concentration of 2 μg/mL just before loading them onto a BD Influx cell sorter. After running negative controls (no YFP, no PI and no YFP, PI only), YFP- positive/PI-negative cells were collected in a low-binding 1.5 mL tube containing 10% FBS in PBS on ice.

Each FACS collection was considered a biological replicate. When possible, at least one male and one female mouse was used per biological replicate to aid doublet identification, and a minimum of two biological replicates were collected per condition. For each replicate, age-matched animals were given the same treatment. See **Supplementary Table 1** for a summary of the experimental design for each sequencing modality in this study.

#### Cell capture and single-cell RNA sequencing

The 10x Genomics droplet-based transcriptome profiling system (Zheng et al Nat Comm 2017) was used to capture single cells, lyse them, and produce cDNA. Reverse transcription (RT) mix was added to the single-cell suspension and loaded onto the Single Cell B Chip. The Chromium Single Cell 3’ GEM, Library, and Gel Bead Kit, Chromium Chip B Single Cell Kit, and Chromium i7 Multiplex Kit were used for RT, cDNA amplification, and library preparation according to the manufacturer’s instructions (Chromium Single Cell 3’ Reagents Kits v2). Indexed single-cell libraries were sequenced in multiplex on Illumina HiSeq 4000 sequencers to produce 100bp paired-end reads.

### scRNA-seq data analysis

#### Initial processing, normalization, and clustering

Fastq files were generated from binary base call files, aligned to *mm10*, quantified, and aggregated using *Cell Ranger* v2.0.0 to produce a feature-barcode matrix containing the number of unique molecular identifiers (UMIs corresponding to cells) associated with a feature corresponding to a gene (row) and a barcode corresponding to a biological sample (column), and a molecule information file containing the number of reads assigned with high confidence to a gene for each UMI. Initial preprocessing of the molecule information file was performed as described in Brann *et al*. (2020). A total of 25,469 cells was initially assayed; after filtering doublets, microvillous cells (176 cells), respiratory epithelial cells (964 cells), and other manually selected cells that were asynchronous with respect to chronological time or separate from the main differentiation process of interest, e.g., mature neurons present at 24h and 48h post-injury, 20,426 cells remained (Brann et al. 2020). Afterwards, for each UMI (cell), the gene expression data were scaled by the median total counts across all cells. Dimensionality reduction was performed using principal component analysis (PCA) and the top 20 principal components were used as input to UMAP (McInnes, Healy, and Melville 2018). Clustering was performed using the *SCANPY* toolkit (Wolf, Angerer, and Theis 2018) using the Leiden algorithm with resolution parameter equal to 1.45, and a manual merging of clusters was performed using known marker genes. We identified 5,418 activated HBCs (HBC*), 7,782 regenerated HBCs (rHBC), 755 globose basal cells (GBC), 2,683 sustentacular cells (Sus), 2,859 immature olfactory sensory neurons (iOSN), and 929 mature olfactory sensory neurons (mOSN).

#### Trajectory inference

The 1,000 most variable genes were selected for dimensionality reduction prior to trajectory inference. PCA on log-transformed counts was performed, calculating the top 25 PCs using the R package *irlba*, which were subsequently reduced to three dimensions using UMAP with parameters min_dist=0.2, n_neighbors=15. Hierarchical clustering based on Euclidean distance in the 3D UMAP-space was performed. The hierarchical tree was cut to obtain nine clusters, which were used as input to *slingshot* for trajectory inference, setting the starting cluster to correspond with HBC* cells.

#### Differential gene expression analysis

Negative binomial generalized additive models (NB-GAM) were fitted using *tradeSeq*. The number of knots for each lineage was set to 6 based on the Akaike Information Criterion (AIC). Top upregulated genes for each lineage, as shown in Figure 1, were identified using the associationTest implemented in *tradeSeq*, requiring a gene to be significant at a 5% nominal FDR level, as well as having a higher estimated average gene expression at the end of the lineage as compared to the lineage starting point based on the NB-GAM.

#### Transcription factor cascade

A list of transcription factors was obtained from the Animal Transcription Factor Database as described previously (Fletcher et al. 2017) and can be found at https://github.com/rufletch/p63-HBC-diff.

We implemented new functionality in tradeSeq to calculate first derivatives of the smooth fitted gene expression functions from the NB-GAM, using finite differencing. Letting *Y*_*gi*_ be the gene expression measure for gene *g* in cell *i*, the tradeSeq model is defined as

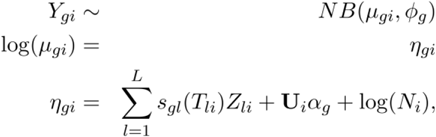

where the mean *μ*_*gi*_ of the negative binomial (NB) distribution is linked to the additive predictor *η*_*gi*_ using a logarithmic link function.

The gene-wise additive predictor consists of lineage-specific smoothing splines *s*_*gl*_, that are functions of pseudotimes *T*_*li*_, for lineages *l ∈* {1,…,*L*}. The binary matrix **Z** = (*Z*_*li*_ ∈ {0,1}: *l* ∈ {1,…,*L*},i ∈ {1,…,*n*}) assigns every cell to a particular lineage based on user-supplied weights. The *n* × *p* matrix **U** is a model matrix allowing the inclusion of known cell-level covariates (e.g., batch, age, or gender), with *i*^th^ row *U*_*i*_ corresponding to the *i*^th^ cell; *α*_*g*_ is a regression parameter vector of dimension *p* × 1.

Differences in sequencing depth or capture efficiency between cells are accounted for by cell- specific offsets *N*_*i*_.

The smoothing spline *s*_*gl*_, for a given gene and *g* lineage *l*, can be represented as a linear combination of *K* cubic basis functions,

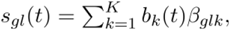

where the cubic basis functions *b*_*k*_(*t*) are enforced to be the same for all genes and lineages. Here, we use six knots, thus, for each gene and each lineage in the trajectory, we estimate *K* = 6 regression coefficients *β*_*glk*_.

#### Estimation of first derivatives

Derivatives of GAMs can be approximated using finite differencing (Wood, 2017). Specifically, assuming a vector of *J* pseudotime grid points 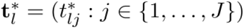 for each lineage *l*, then the derivatives of the splines at these grid points are approximated by

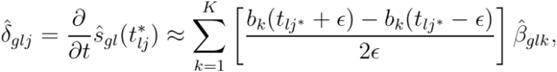

where corresponds to a small finite number, here taken to be 10^−7^.

Standard errors 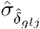 on 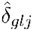 are similarly obtained (Ruppert et al. (2003), Wood (2017)), since the derivatives are a linear combination of the smoother coefficients *β*_*glk*_.

In our application, derivatives for each gene are calculated over a grid *J* = 100 of equally- spaced points for each lineage and, for each grid point, we calculate a thresholded test statistic 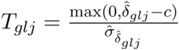, where we set the threshold *c* to be equal to 0.1.

For each gene separately, we considered whether the first derivative of the NB-GAM fit was significantly different from zero for at least one of 100 grid points. Specifically, one-sided *p*- values for the test of the null hypothesis that the derivative is greater than an arbitrary threshold of 0.1 were calculated at each grid point using a standard normal null distribution. Next, (Holm 1979) adjusted *p*-values were computed across grid points and TFs with adjusted *p*-value below a 5% cut-off were declared involved in the cascade. The expression peak of each involved TF was defined as the point where the first derivative crosses zero after the grid point corresponding to the lowest adjusted *p*-value. Note that we merely view the *p*-values as useful numerical summary statistics, without attaching strong probabilistic interpretations to them.

#### Identifying TFs most active in a particular lineage

To identify TFs that are most active in one lineage as compared to other lineages, we start from the set of TFs found to be involved in each lineage from the procedure described in the previous paragraph. Once a TF is included in that set, we consider it to be most active in a particular lineage if the maximum of the estimated lineage-specific smoother is at least 1.5 times larger in that respective lineage as compared to the other two lineages. The 50% increase threshold was chosen to represent a biologically meaningful difference between lineages.

#### Identifying shared TFs across lineages

To identify TFs that are involved across all three lineages, we start from the set of TFs found to be involved in each lineage from the procedure described above. We consider a TF to be ‘shared’, if its expression peak is at a similar pseudotime for all lineages. Out of 90 TFs that peak in all three lineages, we consider 19 TFs to be shared, as the pseudotime difference of their expression peak is lower than 1.

#### Clustering activated HBCs by TF gene expression

Slingshot cluster 8 was used as a proxy for activated HBCs, since it corresponds to the starting point of the trajectory. Gene expression data (counts) for the 19 shared TFs for cluster 8 cells at 24 hpi were scaled, log-normalized, and dimensionally reduced using PCA. The jackstraw procedure within the Seurat toolkit was used to determine the statistical significance of the PCA scores and the elbow method was used to determine the dimensionality of the dataset. The top 15 principal components (PCs) were used as input to UMAP in Seurat for visualization. To cluster the cells, a k-nearest neighbor graph was constructed based on Euclidean distance in the PCA space using the first 7 PCs, and the Louvain algorithm was used to optimize this technique.

#### Clustering and gene set enrichment analysis

Log-transformed and scaled transcription factor expression measures for all TFs found to be involved in each lineage were clustered using hierarchical clustering based on Euclidean distance, and the hierarchical tree was cut at a predefined number of clusters. We somewhat arbitrarily chose 4 clusters for the mOSN lineage and 3 clusters for the sustentacular and rHBC lineages, as more TFs are involved in the nOSN lineage. Gene set enrichment analysis was performed using hypergeometric tests based on the TFs of each cluster using hallmark gene sets from the MSigDB database (Liberzon et al. 2015). For each group, we considered the top 10 enriched gene sets for interpretation.

#### Deconvolution of gene expression to TF activity

For transcription factor activity analysis we only focused on the cells that were assigned to the mOSN lineage, based on the cell assignment procedure from *tradeSeq* and the trajectory fitted using *slingshot* (Street *et al*. 2018, Van den Berge *et al*. 2020). After subsetting, the dataset consist of 14,618 genes and 6,810 cells. The gene regulatory network (GRN) was estimated using grnboost2, implemented in SCENIC v0.10.2 (Aibar et al. 2017). The estimated GRN contains 7,863 genes, 262 transcription factors, and 25,896 edges. The median number of genes regulated by a TF is 27. Transcription factor activity was estimated using *transfactor* (manuscript in preparation), a statistical method which leverages a GRN to assign mRNA molecules to the transcription factors that produced them. The number of molecules produced by each TF in each single cell is then used as a proxy for its activity. Specifically, *transfactor* relies on a hierarchical Poisson model for the number of transcripts produced by each TF for a given gene. The EM algorithm is then used to fit the model and deconvolve TF-specific gene expression from overall gene expression for each gene.

#### HBC dataset integration

To determine whether regenerated HBCs are transcriptionally indistinguishable from resting HBCs, labels for single basal cells (respiratory basal cells, resting HBCs, and activated HBCs) from (Brann et al. 2020) (GSE173999) were transferred to lineage traced HBCs (activated and regenerated) using the IntegrateData function in Seurat.

### Bulk ATAC-seq

#### OE injury, dissociation, and FACS

Injury, removal, and dissociation of the OE was performed as described above with the following changes: wild-type CD1 mice 6–8 weeks-old were used, and only two conditions were assessed -- UI and 24 HPI (3 biological replicates, each consisting of a mix of cells from 2-3 mice, per condition). After papain incubation and 3 washes with 10% FBS in PBS (PBS-FBS), anti-ICAM1-PE antibody was added to the cell suspension to a final concentration of 10 μg/mL and incubated at 4°C for 20 minutes protected from light. Cells were then washed once with ice cold PBS-FBS, resuspended in cold PBS-FBS, and strained through a 35 μm nylon mesh filter cap into a 5 mL polypropylene tube to remove debris. Propidium iodide (PI) was added to cells at a final concentration of 2 μg/mL just before loading them onto a BD inFlux or BD FACSAria cell sorter. After running negative controls (no antibody, no PI and no antibody, PI only), ICAM1-PE-positive/PI-negative cells were collected in a low-binding 1.5 mL tube containing PBS-FBS on ice.

#### Library construction and sequencing

ATAC-seq libraries were generated from FACS-purified ICAM1-PE-positive cells as described in (Corces et al. 2017). Briefly, 10,000–15,000 cells were collected in 100 μl cold PBS-FBS, pelleted at 500 RCF for 5 min at 4°C, and lysed on ice for 3 mins in 50 μl cold lysis buffer (1% Digitonin, 10% Tween-20, 10% NP40 in 1M Tris-HCl pH 7.4, 5M NaCl, 1M MgCl_2_). Cold wash buffer (10% Tween-20 in 1M Tris-HCl pH 7.4, 5M NaCl, 1M MgCl_2_) was added, and tubes were inverted 3 times. Nuclei were then pelleted at 500 RCF for 10 mins at 4°C, supernatant was removed, and 20 μl transposition mix (1X Tagment Buffer, 0.01% Digitonin, 0.1% Tween- 20, 1 μl TDE1) was added to each sample. Nuclei in transposition mix were incubated at 37°C for 30 minutes in a thermomixer with 1000 RPM mixing. Immediately after transposition, samples were placed on ice and 5 volumes of DNA Binding Buffer (Zymo) were added to each. Reactions were cleaned up using a Zymo DNA Clean and Concentrator 5 kit, and samples were eluted in 20 μl Elution Buffer (Zymo).

For library generation, 18 μl of each sample was used as template for 5 cycles of PCR, then qPCR was performed on 4 μl of PCR product to assess the appropriate number of additional PCR cycles. The remaining 36 μl PCR reaction was run for the determined number of cycles. AMPure XP beads were used to select fragments between 180 and 1130bp, DNA concentration was measured using a Qubit dsDNA HS kit, and library quality was visualized using a Bioanalyzer. If fragments less than 100bp were observed, the library was further cleaned up using a Zymo DNA Clean and Concentrator 5 kit and eluted with 10 μl of 10 mM Tris-HCl, pH 8.0. Each replicate per condition was sequenced on a HiSeq 4000 (Illumina) in a different batch such that each sequencing batch contained one replicate from each condition.

### Bulk ATAC-seq data analysis

#### Peak calling

Adapters were trimmed using Trimmomatic (Bolger, Lohse, and Usadel 2014), and reads were aligned to mm10 with bowtie2 (Langmead and Salzberg 2012). After removing reads mapped to the mitochondrial genome, duplicated reads, and reads mapping to any region with fewer than 10 reads with SAMtools (Danecek et al. 2021), the remaining reads were used for peak calling with MACS2 (--shift -50 --extsize 100) (Zhang et al. 2008) and Genrich (https://github.com/jsh58/Genrich), and the intersection of replicate peaks from the two tools were used for differential accessibility analysis (see below).

#### Differential accessibility

Within each condition, replicate peaks identified using a single tool (MACS2 or Genrich) were intersected using bedtools *intersect* (Quinlan and Hall 2010), *then peaks identified using MACS2 were intersected with peaks identified using Genrich using bedtools intersect*. 65,998 total peaks in all samples were quantified using bedtools *merge*. Peaks were then annotated and ATAC-seq signals were quantified and normalized to library size, peak width, and GC-content using *cqn* (Hansen, Irizarry, and Wu 2012). *Differentially accessible (DA) peaks between UI HBCs and 24HPI HBCs were identified using edgeR* (Robinson, McCarthy, and *Smyth 2010) for a total of 15,815 DA peaks*.

### Bulk RNA-seq

#### OE injury, dissociation, and FACS

Injury, removal, dissociation, and FACS were performed as for bulk ATAC-seq, for a total of 2 biological replicates per condition. Each biological replicate consisted of a mix of cells from 2 male and 2 female age-matched mice.

#### Library construction and sequencing

RNA-seq libraries were generated from 20,000–50,000 FACS-purified ICAM1-PE-positive cells collected in 100 μl cold PBS-FBS, pelleted at 500 RCF for 5 min at 4°C, and resuspended in 100 μl TRI Reagent (Zymo). Total RNA was extracted using a Zymo Direct-zol RNA MicroPrep kit, rRNA was depleted using a NEBNext rRNA Depletion Kit (New England BioLabs Cat# E6310), and libraries were made using a NEBNext Ultra II Directional RNA Library Prep Kit for Illumina (New England BioLabs Cat# E7760). All four samples (two biological replicates from two conditions) were sequenced together in one run on a HiSeq 4000 (Illumina).

#### Bulk RNA-seq data analysis

Sequencing reads were mapped to *mm10* using *STAR* v 2.7.1a (Dobin et al. 2013). After removal of duplicated and low-quality reads using SAMtools (Danecek et al. 2021), reads were quantified using HOMER (Heinz et al. 2010). Differential expression was determined using *DESeq2* (Love, Huber, and Anders 2014).

### scATAC-seq with lineage tracing

#### OE dissociation and FACS Purification

Injury, removal, and dissociation of the OE were performed as described above with the following changes: *Krt5-CreER*; *Rosa26*^*eYFP/eYFP*^ mice were injected intraperitoneally once with tamoxifen (0.25 mg tamoxifen/g body weight) at 68 weeks of age, injected with methimazole (50 μg/g body weight, IP) or saline 3 days after tamoxifen administration, and sacrificed 24 h after injury (3 biological replicates per condition, and each replicate consisted of 2–3 mice.

#### Nuclei Isolation

Viable YFP+ cells (range: 900-3600) were sorted into 200 μl of PBS-FBS using a FACSAria (BD), centrifuged at 500g for 5 min at 4°C. After supernatant removal, cells were resuspended in 200 μl 0.04% BSA in PBS and centrifuged again at 500g for 5 min at 4°C. 195 μl supernatant was removed, and 45 μl cold lysis buffer (10mM Tris-HCl pH 7.4, 10mM NaCl, 3 mM MgCl2, 0.1% Tween-20, 0.1% NP40, 0.01% Digitonin, 1% BSA) was added and mixed with the pellet by gently pipetting up and down 3 times. Samples were incubated on ice for 3 min, and 50 μl cold wash buffer (10mM Tris-HCl pH 7.4, 10mM NaCl, 3mM MgCl2, 0.1% Tween-20, 1% BSA) was added to each without mixing. Samples were centrifuged at 500g for 5 min at 4°C. 95 ul supernatant was carefully removed, 45ul cold 1X Nuclei Buffer was added to each sample without mixing, and samples were centrifuged again at 500g for 5 min at 4°C. 48 μl supernatant was removed without disturbing the pellet, which was then resuspended in the remaining (∼5 μl) buffer for use in single cell ATAC-seq.

#### 10x Genomics ATAC-seq library construction

Nuclei isolated as described above were used for preparing single-cell ATAC-seq libraries using a Chromium Single Cell ATAC Library & Gel Bead Kit (PN-1000111) according to the manufacturer’s instructions (10x Chromium Single Cell ATAC Reagents Kits User Guide). Briefly, 5 μl of each sample resuspended in 1X Nuclei Buffer was added to 10 μl Transposition Mix, mixed by pipetting 6 times, and incubated in a thermal cycler at 37°C for 1 hour. After addition of barcoding master mix, the Chromium Chip E Single Cell Kit (PN-1000086) and Chromium i7 Multiplex Kit N, Set A (PN-1000084) were used for GEM generation and library construction. Indexed single-cell libraries were sequenced in multiplex on a NovaSeq (Illumina) to produce 150bp paired-end reads.

### scATAC-seq data analysis

Raw data were processed using the *Cell Ranger* scATAC-seq processing pipeline. Downstream analysis was performed using *ArchR* (Granja et al. 2021) *v0*.*9*.*5, discarding cells that had fewer than 1,000 fragments or a transcription start site (TSS) enrichment score less than 4. Doublets were removed by ArchR* using default settings. Gene activity scores were calculated using default settings in *ArchR*. Latent semantic indexing was performed to obtain 30 reduced dimensions, which were subsequently corrected for sample/batch effects using *Harmony* (Korsunsky et al. 2019). *Clustering was performed on the Harmony* embeddings using the ‘Seurat’ method implemented in *ArchR*, obtaining 10 clusters which were merged manually. For visualization purposes, the Harmony embeddings were used as input to UMAP (McInnes, Healy, and Melville 2018) to reduce it to two dimensions. Gene markers for the HBC cell states were obtained based on the estimated gene activity scores using the getMarkers implementation in *ArchR*, using a Wilcoxon test with a log-fold-change cut-off of 1.25 and Benjamini-Hochberg FDR (Benjamini and Hochberg 1995) nominal level of 1% as thresholds.

Pseudobulking, where counts are summed across cells that share the same cell state/type, was performed using the addGroupCoverages implementation in *ArchR*, creating a minimum of 2 and maximum of 6 pseudobulks for each of the three HBC cell states. Peak calling was performed using *MACS2* (Zhang et al. 2008) on the pseudobulks, resulting in 183,712 peaks. Peak markers for the HBC cell states were obtained as described in the previous paragraph, using a Wilcoxon test statistic with a log-fold-change cut-off of 1.25 and Benjamini-Hochberg FDR nominal level of 1% as thresholds. Motif enrichment as implemented in ArchR (Granja et al. 2021) for the marker peaks of each cell state was performed using the cisbp database (Weirauch et al. 2014).

### Integration of scRNA-seq and scATAC-seq data

*We used Seurat* v3.1.5 for integration of the scRNA-seq and scATAC-seq datasets (Stuart et al. 2019). For the scATAC-seq data, we used the peak matrix and gene activity matrix obtained using *ArchR* (Granja et al. 2021) as described in the previous section. Integration was performed by identifying anchor cells across datasets using canonical correlation analysis on the scRNA-seq gene expression and scATAC-seq gene activity scores. Cell labels were transferred from the scRNA-seq to scATAC-seq data using these anchor cells, as implemented in *Seurat*’s TransferData function. Marker peaks/genes and motif enrichment was performed for each cell identity using *ArchR* as described in the previous section.

### Identification of genes associated with hybrid-specific ATAC-seq peaks

To associate peaks with genes, hybrid-specific marker peaks were saved in a .bed file, which was then entered into GREAT version 4.0.4 using default basal + extension parameters (constitutive 5.0 kb upstream and 1.0 kb downstream, up to 1000.0 kb max extension) with curated regulatory domains (McLean et al. 2010). Genes were ordered by binomial rank and the top 100 significant by region-based binomial FDR Q-value (*Q-val* < 0.05) were considered for validation.

### Histology

#### In-situ hybridization and immunohistochemistry

Nasal tissue was prepared for RNAscope (ACDBio) multiplex fluorescent in-situ hybridization (FISH) from two C57Bl/6J mice at each post-injury time point and one mouse which was uninjured. Briefly, skin was removed from the skull and the bottom jaw and incisors were removed with a razor blade. Nasal cavities were flushed with 4% paraformaldehyde (PFA) via the palate. Heads were then drop-fixed in 4% PFA for 16-18 hours, decalcified in 10% ethylenediaminetetraacetic acid (EDTA) for 3 to 5 days and equilibrated in 30% sucrose in PBS at 4°C before freezing in OCT medium (Fisher Scientific). Sections were sliced to 15µM in a cryostat (Leica Biosystems). Multiplex FISH was performed according to the manufacturer’s instructions (multiplex V2 kit for fixed frozen tissue), with slight modifications: we limited antigen retrieval to 5 minutes and protease III treatment to 10 minutes, except in the case of Figure 2e (uninjured), where the protocol was followed entirely according to the manufacturer’s instructions.

Following FISH, sections were blocked with PBS + 0.1% Tween-20 containing 10% Normal Goat Serum (NGS) for 2 hours and then incubated with an antibody to Keratin 5 (SAB4501651 raised in rabbit from Sigma Aldrich) at 1:500 dilution in PBS + 0.1% Tween-20 containing 1% NGS over the weekend. Sections were then washed in PBS and incubated for 2 hours with a goat anti-rabbit IgG (H+L) highly cross-adsorbed secondary antibody conjugated to Alexa Fluor Plus 647 (ThermoFisher Scientific) at 1:3000 dilution in PBS + 0.1% Tween-20 with 1% NGS. After washing, sections were mounted with glass coverslips using ProLong Gold Antifade Mountant (Invitrogen), and sealed with clear nail polish. Sections and z-stacks were imaged at 20x magnification on a Stellaris 8 confocal microscope (Leica Microsystems).

## Data and code availability

scRNA-seq data have been deposited on the Gene Expression Omnibus (GEO) under the accession number GSE157068 (https://www.ncbi.nlm.nih.gov/geo/query/acc.cgi?acc=GSE157068). scATAC-seq have been deposited on GEO under the accession number GSE246167 (https://www.ncbi.nlm.nih.gov/geo/query/acc.cgi?acc=GSE246167).

Code to reproduce analyses and figures is available on GitHub at https://github.com/koenvandenberge/hbcRegenOE. The code for TF activity analysis will be released once the R package ‘transfactor’ implementing the statistical methodology has been made available.

