## Supplementary Figures for "A Latent Activated Olfactory Stem Cell State Revealed by Single-Cell Transcriptomic and Epigenomic Profiling"

### Supplementary Material for Van den Berge et al.

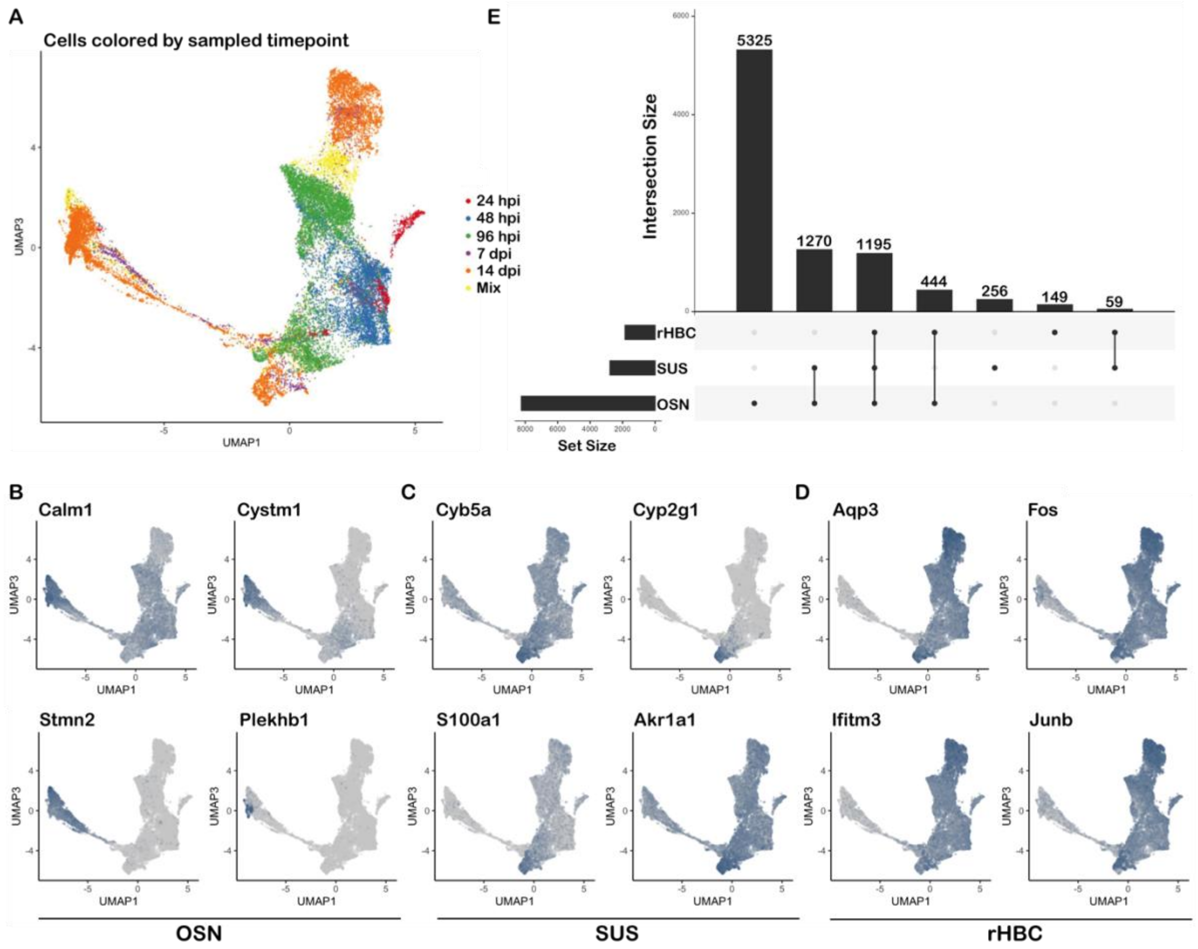


***Supplementary Figure 1**: **(A)** UMAP dimensionality reduction of the cells colored according to their experimentally sampled timepoint. For improved visualization, UMAP dimensions 1 and 3 are visualized. **(B-D)** Markers identified by tradeSeq in 2D UMAP space for the (B) neuronal lineage, (C) sustentaclular lineage, and (D) rHBC lineage. Darker color corresponds with higher expression. **(E)** UpSet plot comparing overlap of differentially expressed genes for each lineage. Left panel: Barplot of the number of DE genes for each lineage on a 5% nominal FDR level. Top panel: Barplot of the number of DE genes in common for every combination of lineages. Center panel: Combination of methods under consideration for the top panel. For example, the first column shows that 5,325 DE genes are only found to be associated with pseudotime in the neuronal lineage.*

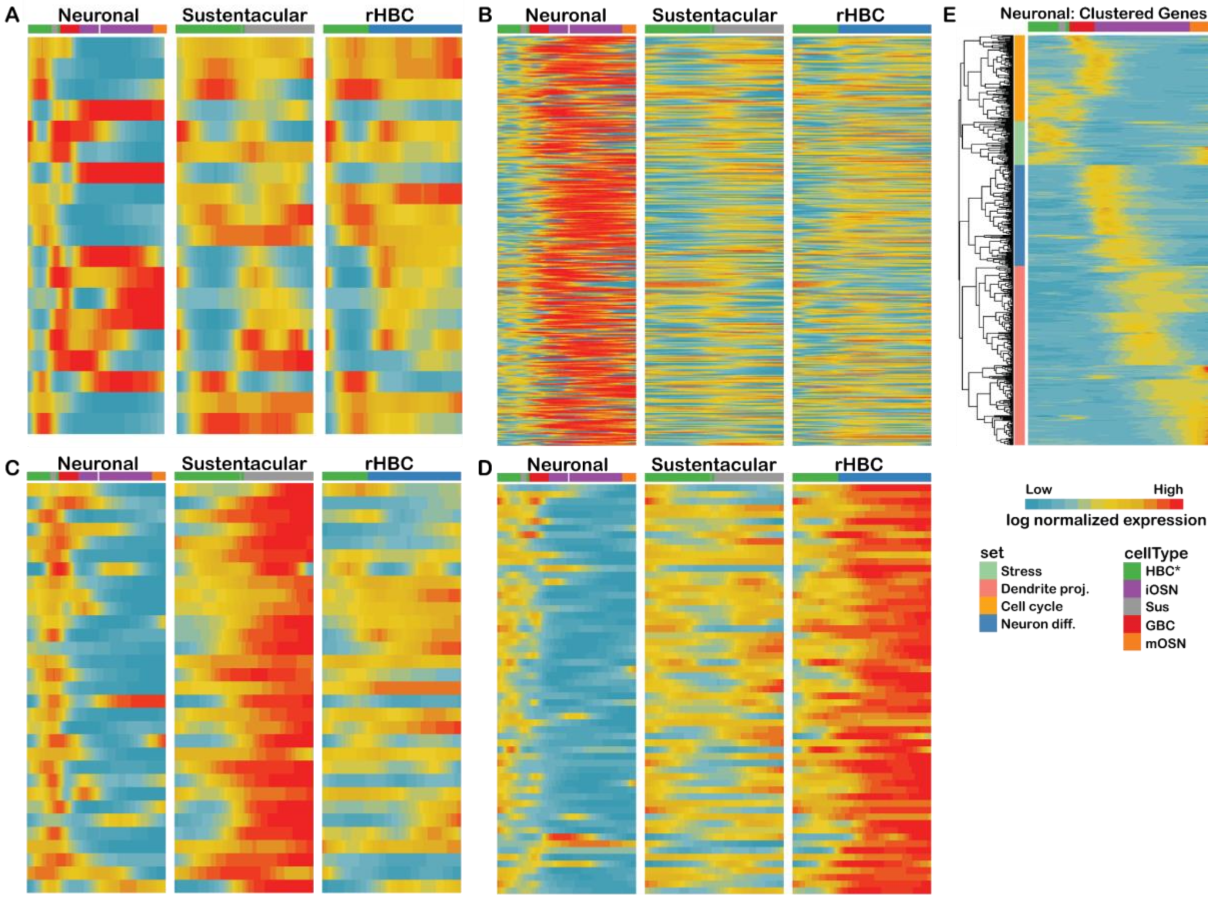


***Supplementary Figure 2****:* ***(A-D)*** *Interpretation of TF expression cascades for shared and lineage specific TFs. Heatmaps for (A) TFs found to be shared across all lineages, or (B) most active in the neuronal lineage, or (C) most active in the sustentacular lineage, or (D) most active in the regenerated HBC lineage. The x-axis for each panel represents 100 bins of pseudotime, and the most abundant cell type in each bin is indicated using the colors at the top of each heatmap. If there are too few cells in a bin, no color is provided. Each row in each heatmap represents scaled log-transformed expression of a transcription factor, normalized to zero mean and unit variance across all lineages. The list of shared TFs is available in Supplementary File 1.* ***(E)*** *Clustered heatmap of scaled log-normalized expression for TFs (rows) involved in the neuronal lineage. The x-axis represents pseudotime and the dominant cell type in each pseudotime bin is indicated at the top of the heatmap. The TFs are clustered into four groups using hierarchical clustering. For each cluster, gene set enrichment analysis is performed; enriched gene sets correspond to known biology and are available in Supplemental File 1.*

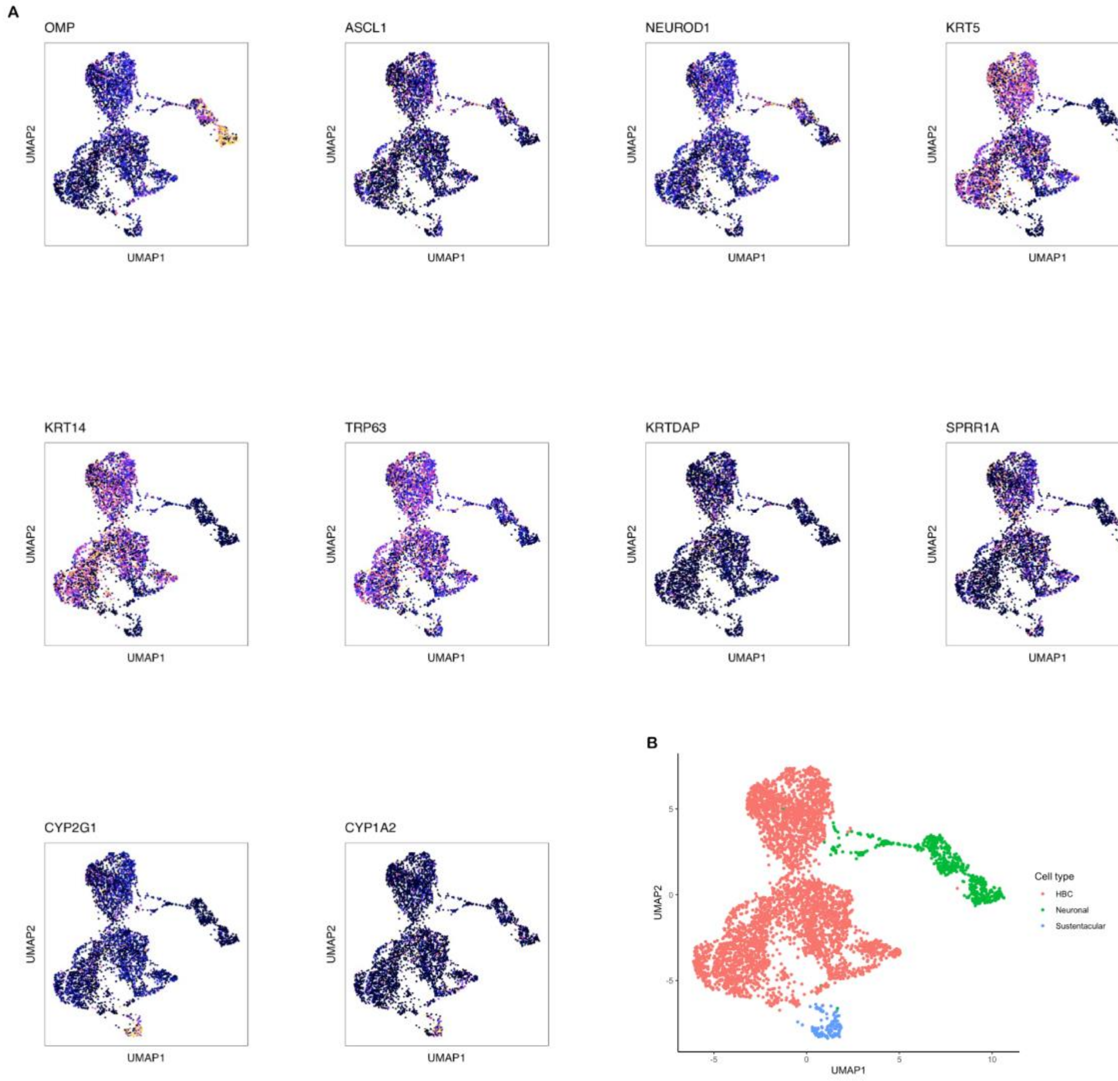


***Supplementary Figure 3****: UMAP dimensionality reduction of all scATAC-seq cells after quality control.* ***(A)*** *scATAC-seq data in 2D UMAP space, colored by known markers. Each panel shows a two-dimensional UMAP visualization of the scATAC-seq cells. Cells are colored according to the estimated gene activity of the corresponding marker gene as indicated by the panel title. Lighter color/yellow corresponds with higher gene activity.* ***(B)*** *UMAP visualization of scATAC-seq data colored according to cell type, as defined by manually merged clusters, with the majority of cells corresponding with the HBC cell type, and two smaller clusters corresponding with neuronal and sustentacular cells.*

A

| | Activated | | Hybrid | | Resting | |
|---|---|---|---|---|---|---|
| Gene | Median | Mean | Median | Mean | Median | Mean |
| Icam1 | 0 | 0.36 | 0.66 | 0.67 | 0.82 | 0.79 |
| Krt5 | 1.02 | 0.95 | 0.43 | 0.55 | 0.78 | 0.75 |
| Krt6a | 0.34 | 0.53 | 0 | 0.18 | 0 | 0.15 |
| Ecm1 | 0.57 | 0.60 | 0.16 | 0.33 | 0 | 0.19 |
| Sprr1a | 0 | 0.36 | 0 | 0.33 | 0 | 0.17 |
| Emp1 | 0.94 | 0.92 | 1.00 | 0.98 | 0.39 | 0.39 |

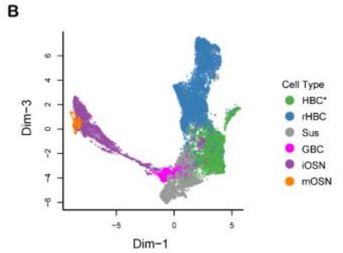


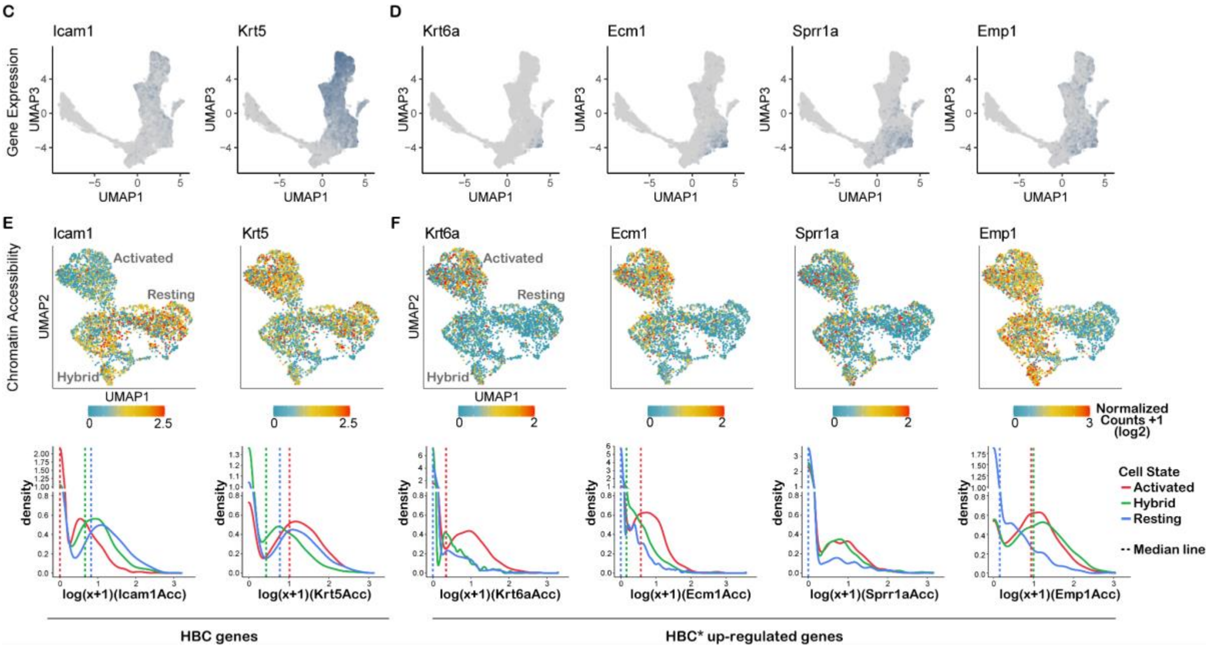


***Supplementary Figure 4**: A subset of wound response genes have similar gene activity in the hybrid and activated states compared with the resting state. (A) Median and mean values of wound response gene activity scores for cells in the scATAC-seq activated, hybrid, or resting clusters. (B) Legend showing cell type clusters by 2D UMAP of scRNA-seq data from lineage traced cells. (C,D) Scaled expression of HBC marker genes (c) and wound response genes (d) visualized by 2D UMAP. Darker color corresponds with higher expression. Wound response genes are enriched in the HBC* cluster. (E) top: 2D UMAP representation of scATAC-seq data where cells are colored according to gene activity scores for the HBC marker gene Icam1 or Krt5; bottom: density plots of gene activity scores for Icam1 or Krt5 in each of the three scATAC-seq clusters representing three HBC states (bottom). (F) UMAP and density plots of gene activity scores for each of four wound response genes.*

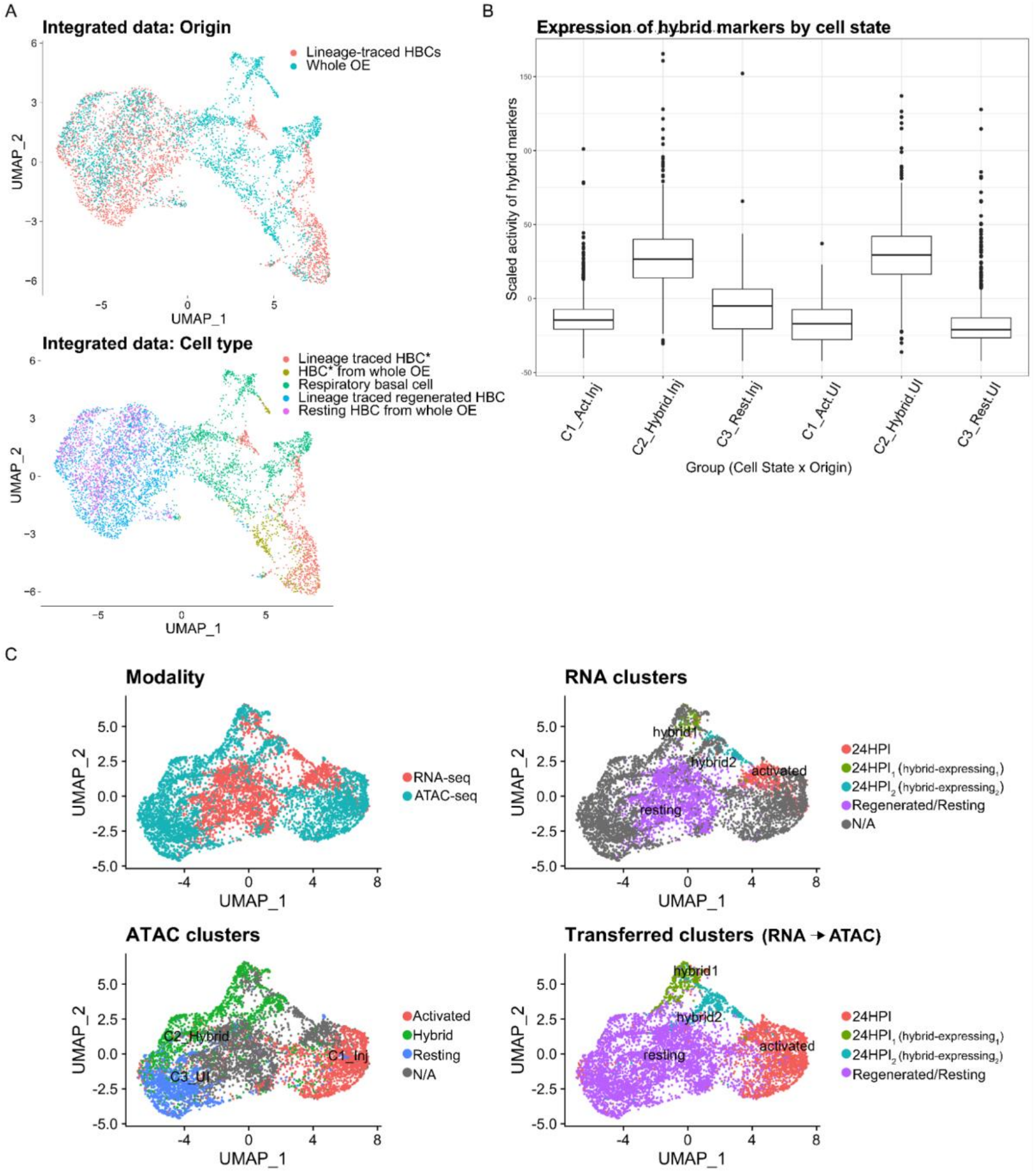


***Supplementary Figure 5:*** *The 'hybrid' epigenomic state represents injured and uninjured cells and is marked by genes that are expressed after injury. (A) Joint dimensionality reduction of transcriptomes of basal cells from lineage-traced OE after injury and whole OE before and after injury shows co-clustering of regenerated and resting HBCs. (B) Scaled gene activity of hybrid markers for injured versus uninjured cells in each scATAC-seq cluster. The x-axis represents the interaction of cell state (C1_Act: activated state, C2_Hybrid: hybrid state and C3_Rest: resting state) with origin (Inj: injured, UI: uninjured). The y-axis represents scaled estimated gene activity scores based on the scATAC-seq data. The activity for all markers of the hybrid cluster are plotted, showing that both the uninjured and the injured cells within the hybrid cluster contribute to the hybrid markers. (C) Joint dimensionality reduction and cell label transfer of RNA-seq to ATAC-seq data reveals '24HPI1' and '24HPI2' RNA-seq subclusters, which express hybrid markers, mapped to the 'hybrid' cluster in the ATAC-seq dataset.*

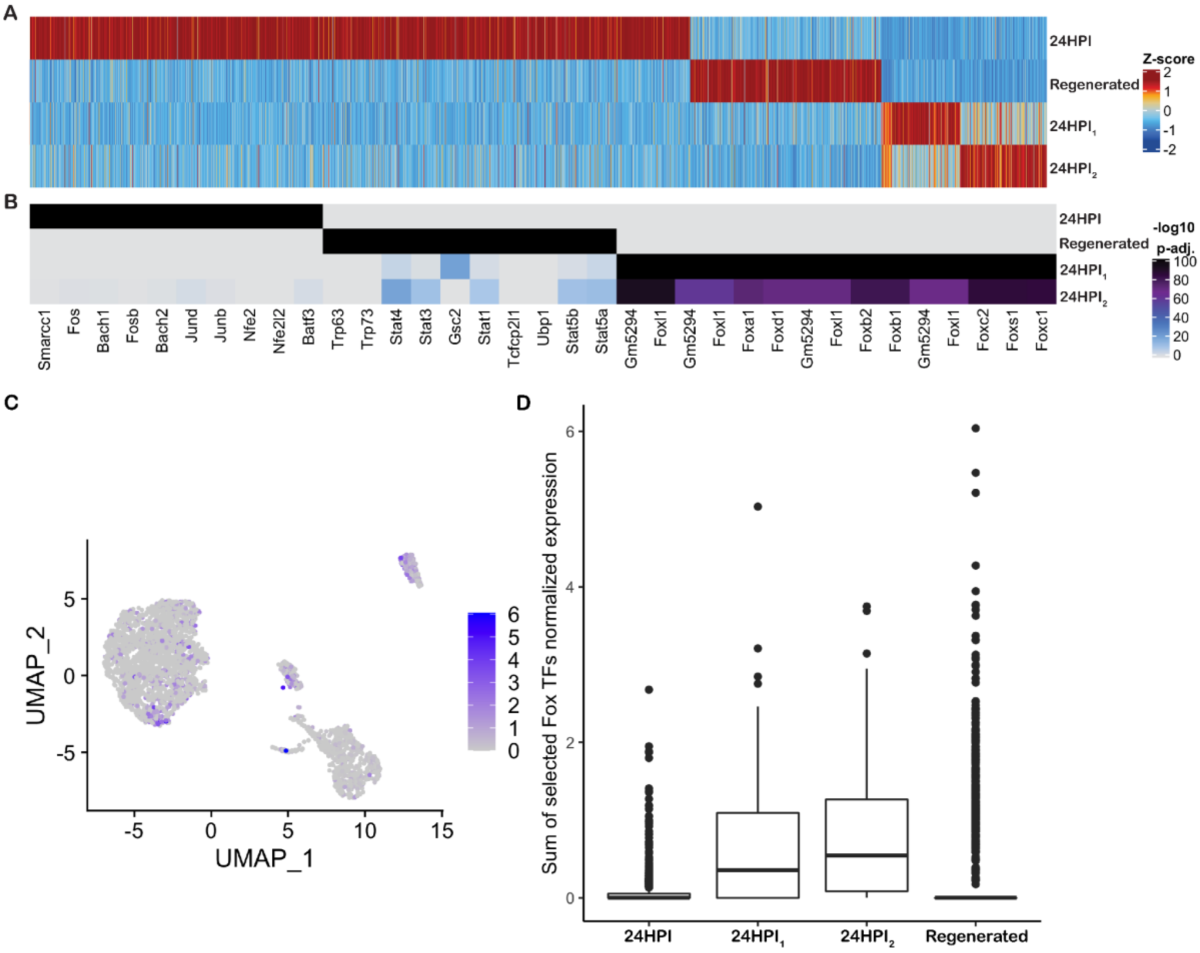


***Supplementary Figure 6***: *Enrichment of Fox transcription factors in the 24HPI₁ and 24HPI₂ subclusters.* ***(A)*** *Heatmap of scaled gene activity scores based on scATAC-seq data, showing markers for 24HPI and regenerated subclusters.* ***(B)*** *Heatmap of adjusted p-values for the top 10 most enriched transcription factor motifs in marker peaks of each of each of the identified scATAC-seq subclusters.* ***(C)*** *Normalized expression of Fox TFs in UMAP space. Darker color corresponds with higher gene expression.* ***(D)*** *Boxplots of the sum of normalized expression of Fox TFs across the four subclusters.*

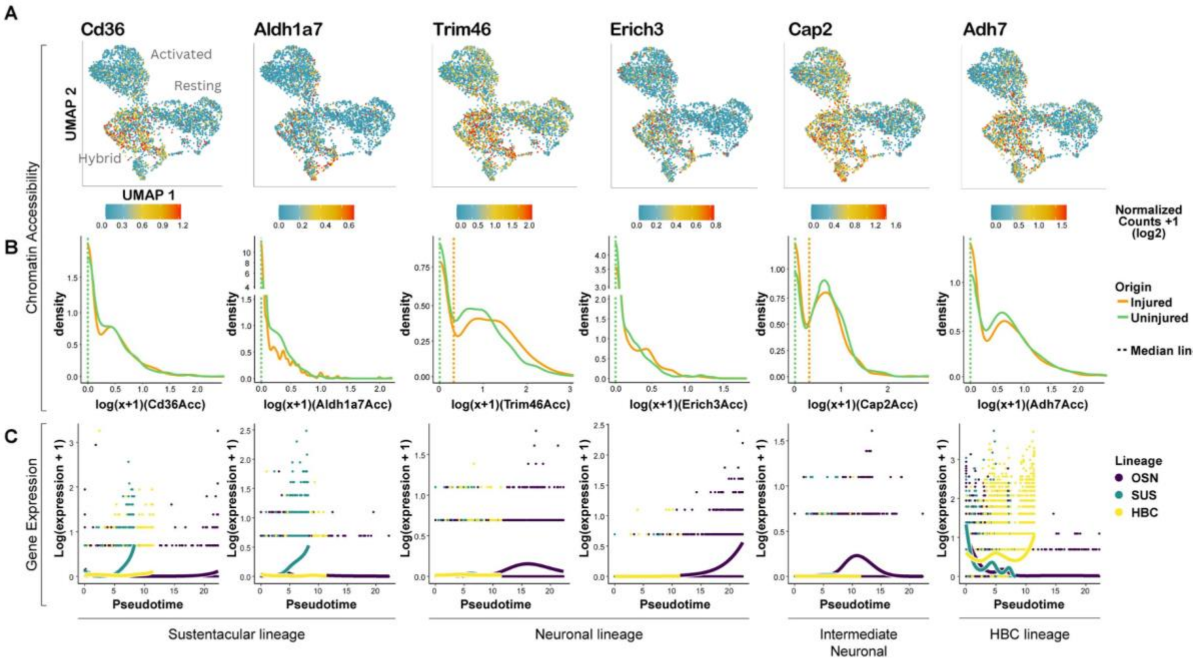


***Supplementary Figure 7****: Lineage-specific genes are accessible in both uninjured and injured hybrid HBCs.* ***(A)*** *UMAP plots of normalized gene activity scores for lineage-specific genes in resting, hybrid, and activated HBCs.* ***(B)*** *Density plots of gene activity scores for each gene for hybrid cluster cells labeled by treatment.* ***(C)*** *Expression of each gene in each lineage over pseudotime.*

| Modality | FACS Purification Strategy | Conditions | Biological replicates per condition | Mice pooled per bio. replicate | Age range |
|---|---|---|---|---|---|
| scRNA-seq | Krt5CreER; R26-YFP lineage traced cells | 24, 48, 96 HPI 7dpi, 14dpi | 2 | 1-6 | 3-7 weeks |
| bulk RNA-seq | ICAM+ HBCs | UI, 24 HPI | 2 | 4 (2M, 2F) | 6.1-8 weeks |
| scATAC-seq | Krt5CreER; R26-YFP lineage traced cells | UI, 24 HPI | 3 | 3 | 5.6-7.9 weeks |
| bulk ATAC-seq | ICAM+ HBCs | UI, 24 HPI | 4 | 2-3 | 6-8 weeks |

***Supplementary Table 1****: Summary of purification strategies, conditions, biological replicates, and age ranges of animals used in this study for each sequencing modality.*
